# Dynamics of GLP-1R peptide agonist engagement are correlated with kinetics of G protein activation

**DOI:** 10.1101/2021.03.10.434902

**Authors:** Giuseppe Deganutti, Yi-Lynn Liang, Xin Zhang, Maryam Khoshouei, Lachlan Clydesdale, Matthew J. Belousoff, Hari Venugopal, Tin T. Truong, Alisa Glukhova, Andrew N. Keller, Karen J. Gregory, Katie Leach, Arthur Christopoulos, Radostin Danev, Christopher A. Reynolds, Peishen Zhao, Patrick M. Sexton, Denise Wootten

## Abstract

The glucagon-like peptide-1 receptor (GLP-1R) has broad physiological roles and is a validated target for treatment of metabolic disorders. Despite recent advances in GLP-1R structure elucidation, detailed mechanistic understanding of how different peptides generate profound differences in G protein-mediated signalling is still lacking. Here we have combined cryo-electron microscopy, molecular dynamics simulations, receptor mutagenesis and pharmacological assays, to interrogate the mechanism and consequences of GLP-1R binding by four peptide agonists; glucagon-like peptide-1, oxyntomodulin, exendin-4 and exendin-P5. These data revealed that distinctions in peptide N-terminal interactions and dynamics with the GLP-1R transmembrane domain are reciprocally associated with differences in the allosteric coupling to G proteins. In particular, transient interactions with residues at the base of the binding cavity correlate with enhanced kinetics for G protein activation, providing a rationale for differences in G protein-mediated signalling efficacy from distinct agonists.

## INTRODUCTION

The glucagon-like peptide-1 receptor (GLP-1R) is widely expressed in many tissues and mediates the action of the gastrointestinal peptide hormone, glucagon-like peptide-1 (GLP-1)^1^. GLP-1 mediates numerous physiological effects that are desirable in the management of type 2 diabetes and obesity, including regulation of insulin secretion, slowing gastric emptying, suppressing appetite and regulating carbohydrate metabolism. Numerous endogenous agonists activate the GLP-1R, including several forms of GLP-1, oxyntomodulin and glucagon, and multiple exogenous peptide agonists are approved, or in clinical development, for the treatment of type 2 diabetes and/or obesity^1, 2^. However, these have different therapeutic efficacies for glucose control, weight loss, improved cardiovascular outcomes, as well as side effect profiles, such as nausea and vomiting^2^. These differential effects may be attributed to their pharmacokinetic profiles and/or distinctions in how each peptide binds and activates the GLP-1R.

The GLP-1R is a class B1 G protein-coupled receptor (GPCR) that mediates its effects via coupling to heterotrimeric G proteins^1^. The receptor predominantly couples to the stimulatory G protein G_s_ to raise cAMP levels within the cell, however it pleiotropically couples to multiple transducers, including other G protein subtypes and regulatory proteins^1, 3^. When compared to GLP-1, other GLP-1R agonists can display differential efficacies within a single signalling pathway, as well as preferential signalling towards individual pathways at the expense of others^4–8^. These phenomena lead to biased agonism, which is commonly observed when GLP-1R agonists are assessed across multiple signalling pathways. However, the molecular basis for how individual agonists can promote profound differences in pharmacology is still poorly understood.

Class B1 GPCRs bind their peptide agonists via a two-domain model, whereby the C-terminus of the peptide interacts with the receptor extracellular N-terminal domain (ECD) promoting an “affinity trap” that enables the engagement of the N-terminus of the peptide with the receptor transmembrane domain (TMD), with interactions with the TMD required for receptor activation ^9^. In recent years, advances in cryo-electron microscopy (cryo-EM) have enabled structural determination of a large number of class B1 GPCRs bound to their endogenous agonists, and coupled to G_s_, including that of the GLP-1R, which confirm engagement of these peptides with both the ECD and the TMD^10–21^.

Naturally occurring GLP-1R peptide agonists share a conserved N-terminal sequence with GLP-1, including oxyntomodulin and the first FDA approved GLP-1R agonist, exendin-4 (Figure 1A). Despite this conservation, these peptides induce distinct signalling profiles and have different mechanisms for receptor interaction^4^. Truncation of just two N-terminal residues of GLP-1 decreases affinity by 100-300-fold and potency for cAMP signalling by 1,000-10,000-fold^22^. In contrast, truncation of the first two residues of exendin-4 does not significantly alter its affinity, but does lower potency for cAMP signalling by approximately 200-fold ^23–25^. Unlike naturally occurring GLP-1R agonists, exendin-P5 contains a unique N-terminal sequence (Figure 1A). While this peptide is a potent agonist for cAMP production, it is a biased agonist, with preference for G protein-mediated signalling relative to *β*-arrestin recruitment when compared to GLP-1 and exendin-4^26^. Intriguingly, this peptide also displays a unique *in vivo* profile relative to exendin-4 with improved ability to reduce hyperglycaemia in animal models of diabetes.

**Figure 1.**
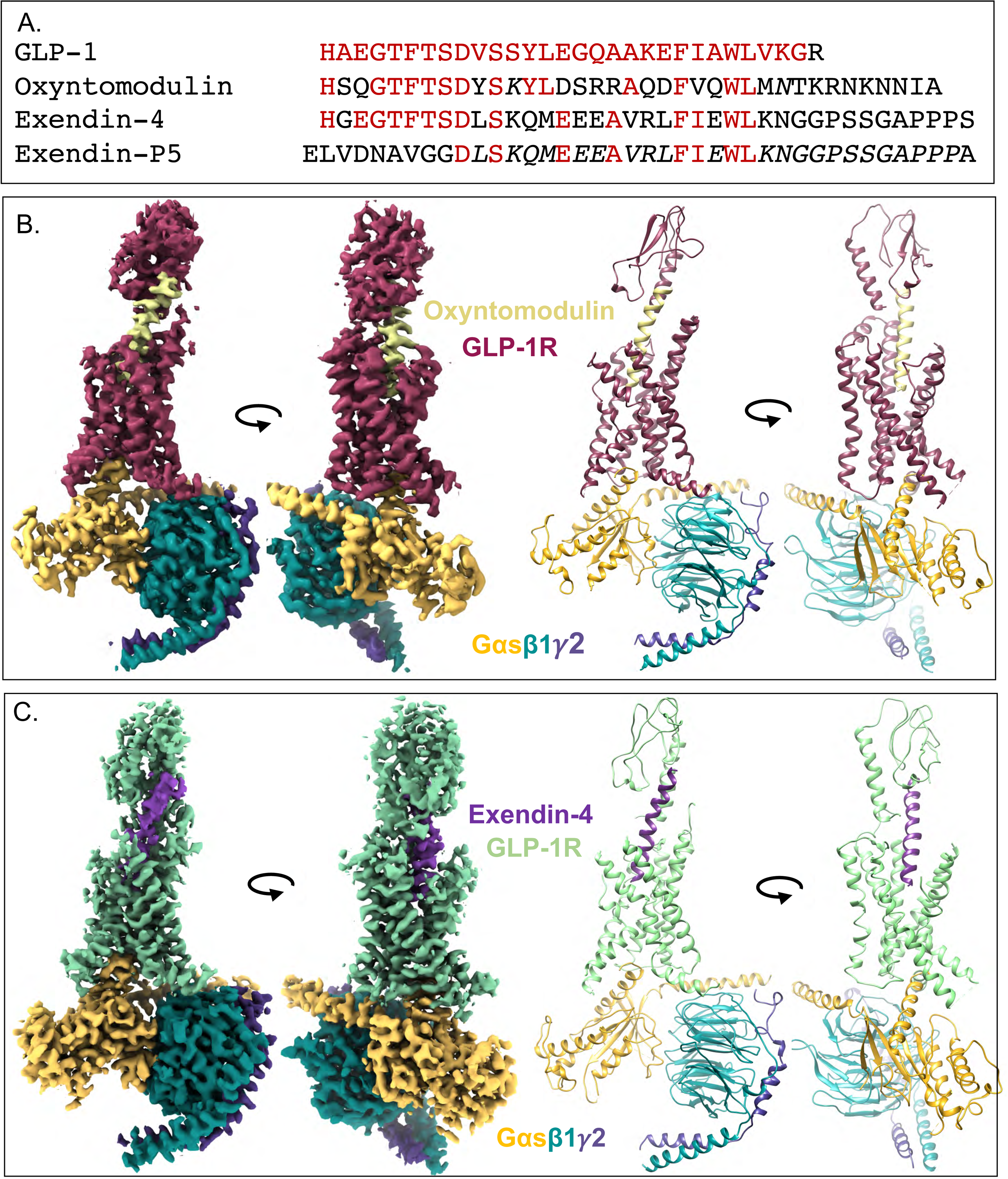
Cryo-EM structures of GLP-1R:G_s_ complexes with different agonists. **A.** Sequences of peptides assessed in this study. **B-C,** orthogonal views of the cryo-EM maps (left), and the backbone models built into the maps in ribbon format (right) for oxyntomodulin (B) and exendin-4 (C) bound GLP-1R:G_s_ complexes. Colouring denotes the protein segments as highlighted on the figure panels

Due to the therapeutic implications of biased agonism and differential efficacy, understanding the molecular details of how different peptides engage and activate the GLP-1R is crucial. Structures of the GLP-1R in complex with G_s_, bound with GLP-1 and exendin-P5, as well as non-peptide ligands, have been determined using cryo-EM^13, 16, 27^. Coupled with mutagenesis data, these structures provide initial molecular insights into biased agonism and differential efficacy. The conformation of TM6-ECL3-TM7 has been correlated with the distinct signalling profiles of GLP-1 and exendin-P5, and along with the conformation of TM1, TM2 and ECL2, are key receptor domains important for GLP-1R cAMP signalling, and biased agonism^3, 13, 28, 29^. However, a detailed mechanistic understanding of GLP-1R activation linked to downstream signalling is still lacking, with the existing structural information unable to fully explain how differential peptide efficacies arise, or the differential requirements of the N-terminus and C-terminal sequences of peptide analogues. We therefore sought to investigate the molecular mechanisms by which GLP-1, oxyntomodulin, exendin-4 and exendin-P5 bind and activate the GLP-1R using a combination of structural biology, molecular dynamics simulations and pharmacological studies combined with extensive receptor mutagenesis.

## RESULTS

### Cryo-EM determination of the GLP-1R:G_s_ complex bound by oxyntomodulin and exendin-4

CryoEM structures of exendin-4- and oxyntomodulin-bound GLP-1R-G_s_ complexes were determined using established methodology^12–14^. Purified complexes (Supplemental Figure 1) that contained all expected components were vitrified and imaged by single particle cryo-EM on a 300kV Titan Krios, with (oxyntomodulin) or without (exendin-4) a Volta Phase Plate (VPP). Processing these datasets yielded consensus maps with global resolutions of 3.3 Å (oxyntomodulin) and 3.7 Å (exendin-4) at gold standard FSC 0.143 (Figure 1B-C, Supplemental Figure 1). As there was only limited density for the α-helical domain (AHD) of the Gα subunit, this was masked out during the refinement. Similar to previous peptide-bound:GLP-1R:G_s_ complex structures, the highest resolution was observed within the G protein and receptor TMD, with lower resolution in the extracellular half of the receptors, including the ECD (Supplemental Figure 1), indicative of greater flexibility in these regions.

The cryo-EM map for the oxyntomodulin bound complex enabled robust modelling and confident assignment of most of the side chain rotamers for oxyntomodulin, the G protein and the receptor TMD (Supplemental Figure 2A), with the exception of extracellular loop (ECL) 1 and intracellular loop (ICL) 3, which were not modelled. The ECD was less well resolved, however the protein backbone could be *ab initio* modelled into the density. The exendin-4 bound complex had lower global resolution however, robust modelling into the map could be performed for the majority of the peptide, the G protein and receptor TMD (Supplemental Figure 2B). ECL1, ECL3 and ICL3 were not modelled as the density was less well resolved indicative of higher flexibility within these domains. The low resolution within the ECD of the exendin-4-bound map precluded confident modelling; as such the ECD was rigid body fitted to the density, followed by MD refinement of the backbone.

### General features of peptide bound GLP-1R:G_s_ complexes

The oxyntomodulin and exendin-4 bound GLP-1R:G_s_ complexes exhibited key features of active state class B1 GPCRs (Supplemental Figure 3) and are consistent with the general features of the active state GLP-1R observed previously, when bound by other agonists^13, 16, 27^. Relative to the inactive GLP-1R^30^, this includes an upwards and clockwise rotation of the ECD relative to the TMD, a reorganisation of the extracellular TM regions to accommodate peptide engagement and rearrangement of a conserved central polar network to stabilise a sharp kink within the centre of TM6, which facilitates the large outward movement of this TM at the intracellular face that is required to accommodate G protein binding. Similar to other peptide-bound GLP-1R structures^13, 16^, exendin-4 and oxyntomodulin adopted a continuous alpha helix, with their C-terminus bound within the ECD and their N-terminus bound deep within the TMD forming extensive interactions with residues within TM1, TM2, TM3 TM5, TM6, TM7 and ECL2 (Figures 1-2, Supplemental Figure 4).

**Figure 2.**
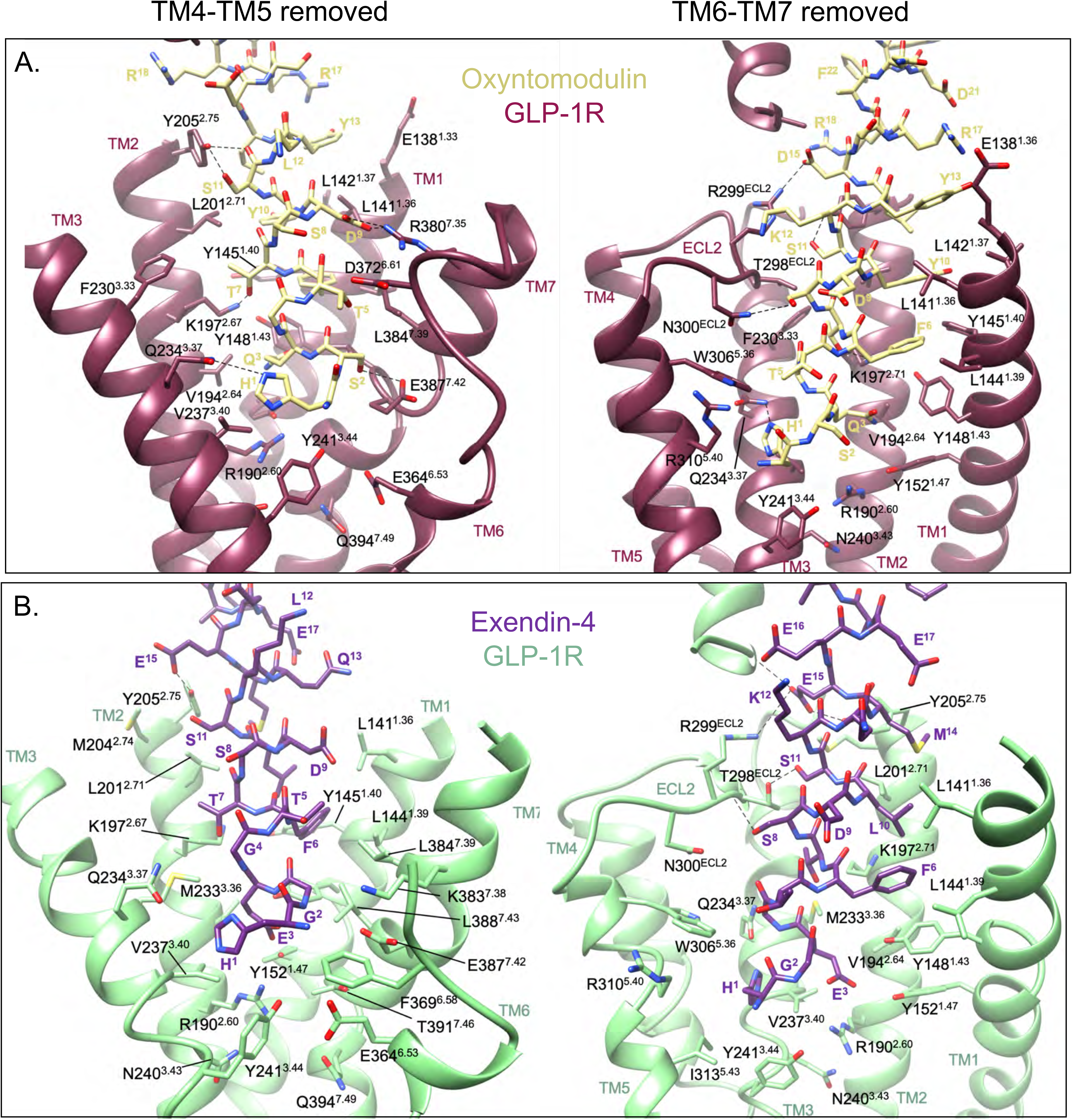
Interactions of oxyntomodulin and exendin-4 peptides within the TMD binding cavity of the GLP-1R. **A**, GLP-1R (dark pink) and oxyntomodulin peptide (pale yellow); **B,** GLP-1R (pale green) and exendin-4 (purple). For each binding site two views are depicted for clarity; Left, side view of the TM bundle viewed from the upper portion of TM4/TM5 where TM4-ECL2-TM5 have been removed; Right; side view of the TM bundle viewed from the upper portion of TM6/TM7 where TM6-ECL3-TM7 have been removed. Dashed lines depict hydrogen bonds as determined using default settings in UCSF chimera. Superscript numbering for receptor residues refers to the generic Wootten et al. class B1 numbering system^38^.

Comparison of the oxyntomodulin-bound structure with the previously determined high-resolution GLP-1-bound GLP-1R (PDB 6X18)^13, 16^ reveals remarkably similar TMD conformations, with only a small difference in position of the ECD relative to the bundle (Figure 3A). In addition, the side chain rotamers within the TMD cavity were also similar, albeit the strength and nature of their interactions with the bound peptides vary (Figure 3B). Exendin-4 also engages the GLP-1R in a comparable manner, with the ECD adopting a similar conformation to the GLP-1-bound receptor (Figure 3A). However, within the TMD, the exendin-4 bound cavity is more open, predominantly due to a more outward location of TM1, however, TM2, TM4-ECL2-TM5 and TM7 are also located further from the centre of the bundle (Figure 3B). Nonetheless, the receptor side chains within the pocket adopt similar rotamers. Comparison of the exendin-4 and exendin-P5 bound (PDB 6B3J) structures revealed similarities in their ECD location and greater similarities of the backbone orientations for TMs 1-5, relative to the GLP-1- and oxyntomodulin-bound complexes (Figure 3). However, the TMD binding cavity is even more open in the presence of exendin-P5, due to a more outward location of TM7. While the top of TM6 and ECL3 could not be confidently modelled for the exendin-4 complex, the portion of TM6 and TM7 that were modelled, along with the weak density corresponding to ECL3 supports a backbone conformation more similar to GLP-1 and oxyntomodulin bound receptors, rather than exendin-P5, albeit it likely that this region is more conformationally dynamic.

**Figure 3.**
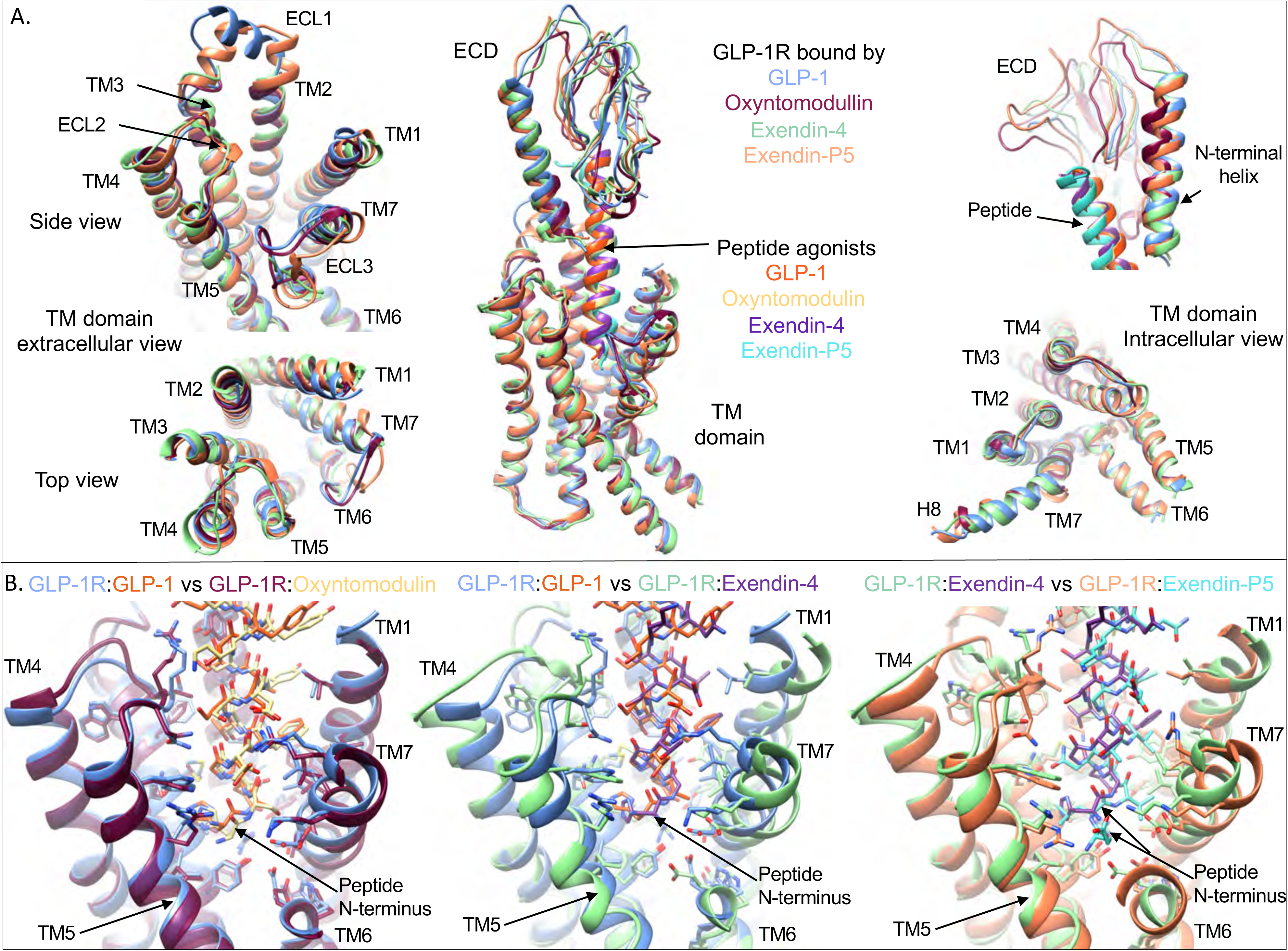
Comparisons of the GLP-1R conformations and binding pockets stabilised by GLP-1, exendin-4, oxyntomodulin and exendin-P5. **A,** Superimposition of the receptor from the GLP-1R:G_s_ complex structures bound with GLP-1 (6X18^16^ – receptor-blue, peptide-orange), oxyntomodulin (receptor-dark pink, peptide-pale yellow), exendin-4 (receptor-pale green, peptide-purple) and exendin-P5 (6B3J^13^ – receptor-pale orange, peptide-cyan). **Middle,** Overlay of full length receptors with bound peptides; **Left**, close up of extracellular portion of the receptor TMD viewed from the side (top) and looking down on the TMD binding cavity (bottom); **Right**, close up of the ECD showing the distinct location of the ECD N-terminal *α*-helix and the location of the peptide C-terminus in the different structures (top) and the receptor TM domain viewed from the intracellular G protein binding site. **B,** Superimposition of the peptide binding sites within the GLP-1R TMD comparing GLP-1 with oxyntomodulin (left), GLP-1 with exendin-4 (middle) and exendin-4 with xendin-P5 (right). Colouring denotes the different peptide bound receptors as highlighted on the figure panels.

### Cryo-EM structures, molecular dynamics and mutagenesis reveal distinct dynamic interactions of individual peptides within the GLP-1R binding site

Atomic modelling into the static consensus cryo-EM maps revealed specific details regarding the interactions of oxyntomodulin and exendin-4 with the GLP-1R. These are reported in Supplemental Table 1 and the interactions of the peptide N-termini with the TMD are shown in Figure 2. The peptide N-terminus is highly conserved between GLP-1, exendin-4 and oxyntomodulin, and as such, a large number of receptor contacts are also conserved, whereas these are more divergent when compared with exendin-P5 (Supplemental Table 1). To interrogate the relative importance of consensus structure interactions for receptor binding and activation, we employed receptor mutagenesis, where each residue within the TMD that formed an interaction with any of the four peptides in the static cryo-EM structures was mutated to alanine (with the exception of A368^6.57^, which was mutated to glycine), and the binding affinity and cAMP signalling of each peptide was assessed. From concentration response curves (Supplemental Figures 5-8), pIC_50_ values, and transduction ratios (log*τ*_c_/K_A,_ where receptor expression was also taken into account) that quantify signalling efficiency, were calculated. These were compared between the mutant and wildtype receptors to assess the impact of the mutation on affinity and signalling of each peptide (Supplemental Figures 9-10, Supplemental Table 2), and these were mapped onto the cryo-EM structures (Figure 4). When comparing the global mutagenesis profile, the effects on exendin-4 were similar to those on GLP-1, with a strong positive correlation observed for the mutagenesis data for both affinity and cAMP production, albeit that the effect on cAMP signalling was generally smaller for exendin-4 (Figures 4-5). Interestingly, while oxyntomodulin-bound GLP-1R displayed a more similar TMD conformation to that bound to GLP-1 in the static cryo-EM structure, the effect of mutagenesis was more divergent, with numerous mutations differentially impacting oxyntomodulin affinity and/or signalling data relative to GLP-1 (Supplemental Figures 9-10, Supplemental Table 2). Nonetheless, there was still a significant weak positive correlation across all mutant datasets and similar to exendin-4, mutations affecting both peptides generally had a greater effect on GLP-1 than oxyntomodulin signalling (Figures 4-5). In contrast, the exendin-P5 mutagenesis profile was very distinct from the other peptides with few mutations altering exendin-P5 affinity and only a very weak, albeit significant, correlation with the effect of mutations on GLP-1 in cAMP signalling assays (Figures 4-5, Supplemental Figures 9-10). In addition, there was no correlation between the oxyntomodulin and exendin-P5 mutagenesis when assessing the effect of mutations as a whole, albeit there were select mutations that exhibited similar effects on the signalling of both peptides (Figures 4-5, Supplemental Figures 9-10).

**Figure 4.**
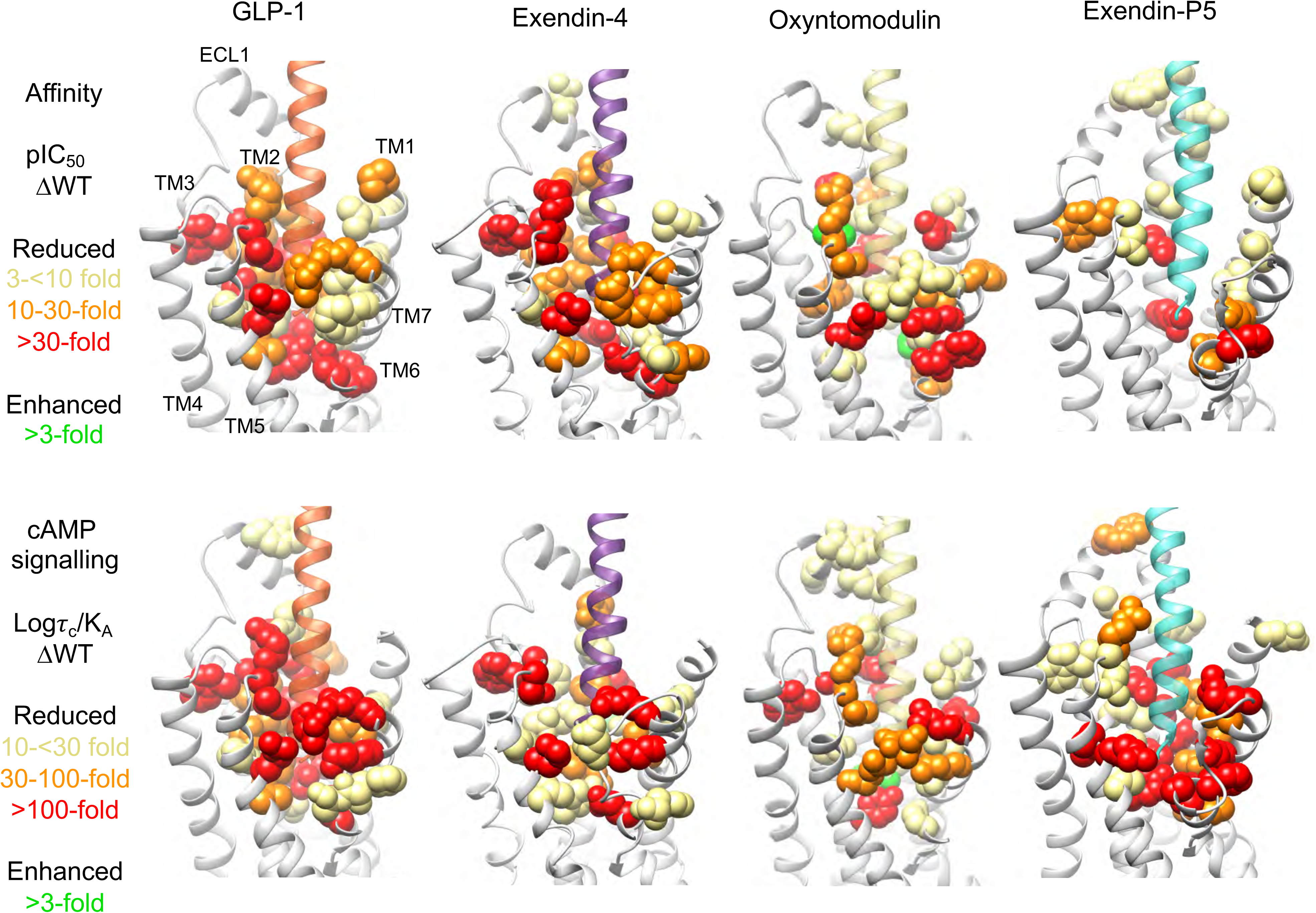
Heat maps depicting the 3D representation the effect of alanine mutation of residues within the TMD peptide binding cavity on affinity and signalling. Models of the peptide bound GLP-1Rs showing residues (in space fill) that altered affinity (top) or signalling (bottom) of GLP-1, exendin-4, oxyntomodulin and exendin-P5 relative to the wild type receptor when mutated. These are coloured depending on their level of effect highlighted in the colour key. TM domains are labelled on the GLP-1-bound model depicting affinity changes, with the same receptor views used for the remaining models.

**Figure 5.**
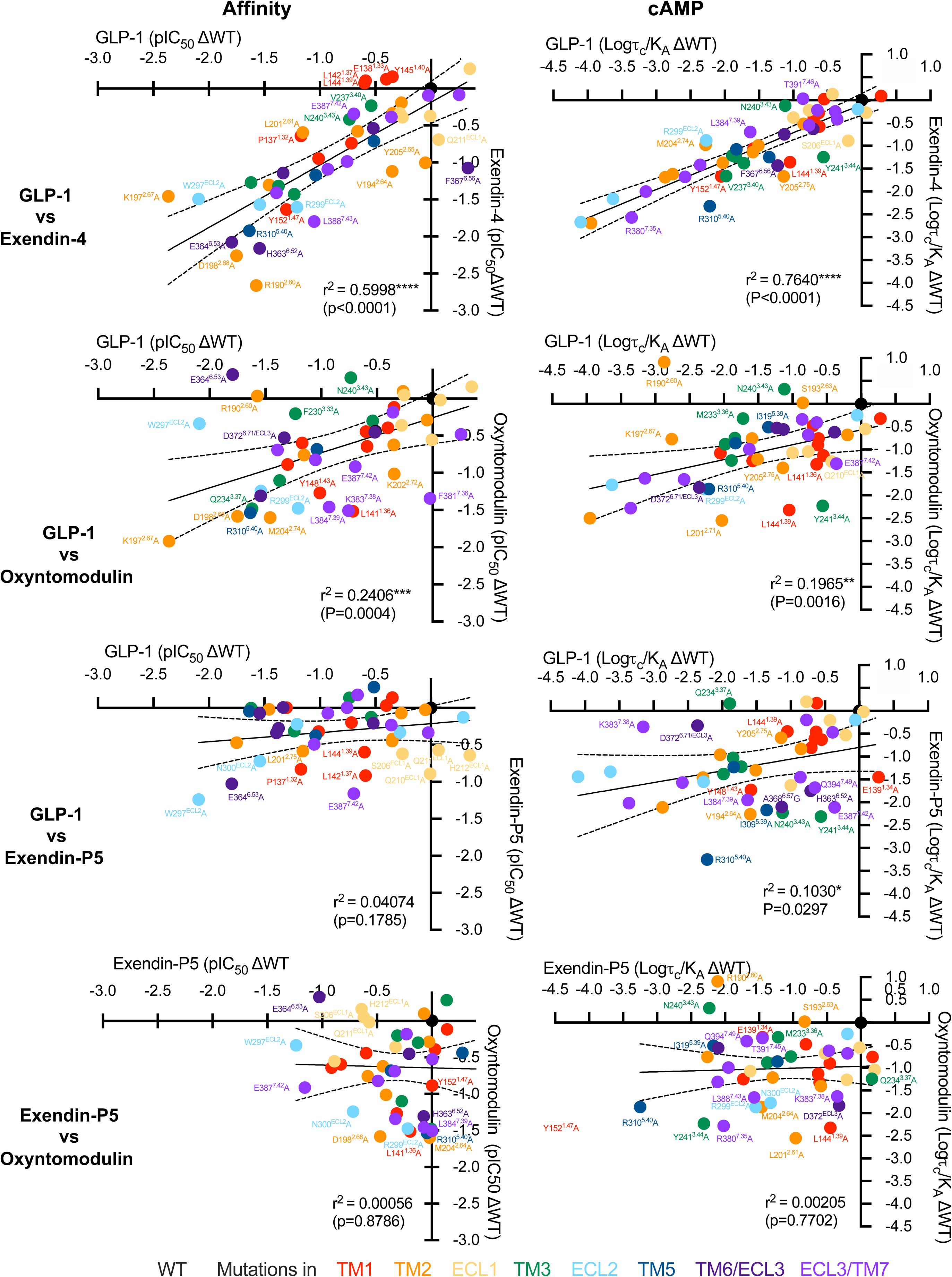
Correlation plots of the changes in peptide affinity and efficacy for TMD mutations relative to the wildtype receptor. **A,** GLP-1 vs exendin-4; **B**, GLP-1 vs oxyntomodulin; **C**, GLP-1 vs Exendin-P5; **D**, Exendin-P5 vs Oxyntomodulin. Data were fit by linear regression and Pearson correlations (r) were determined and squared (r^2^), and statistically assessed to calculate the P value using a two-tailed analysis in Prism 9. The line of regression and 99% confidence intervals are displayed. Mutations are coloured relative to the receptor TM or ECL that they are located as indicated in the legend. Mutant receptors that fall outside of the 99% confidence intervals are labelled.

The divergence in the effect of mutagenesis of residues comprising the TM binding pocket, even where interactions in the static consensus structures were similar, confirmed that static visualisation of complexes is insufficient to fully understand binding and activation mechanisms, and suggests that the dynamics of peptide-receptor engagement likely play a critical role. Therefore, we probed the stability and dynamics of each receptor complex in a simulated POPC lipid environment over microseconds of molecular dynamics (MD) simulations (Supplemental Movie 1). Regions that were not modelled in the cryo-EM maps due to low resolution were first modelled and the receptor complex energy minimised, prior to commencing the simulations. The main peptide-receptor interactions identified in the MD are summarised in Figure 6, Supplemental Figure 11, and Supplemental Table 3. The MD studies revealed that very few residues, particularly within the TMD, formed stable peptide contacts, instead these interactions were transient (Supplemental Figure 11). Nonetheless, there are common residues within the ECD, TM1, TM2, ECL2, TM5, TM6 and TM7 that interact with all four peptides, albeit the nature and stability of these interactions differed considerably between the agonists. In contrast, more divergent interaction patterns were observed within the TMD-ECD linker region (stalk), ECL1, TM2 and ECL3/TM7 (Supplemental Figure 11).

**Figure 6.**
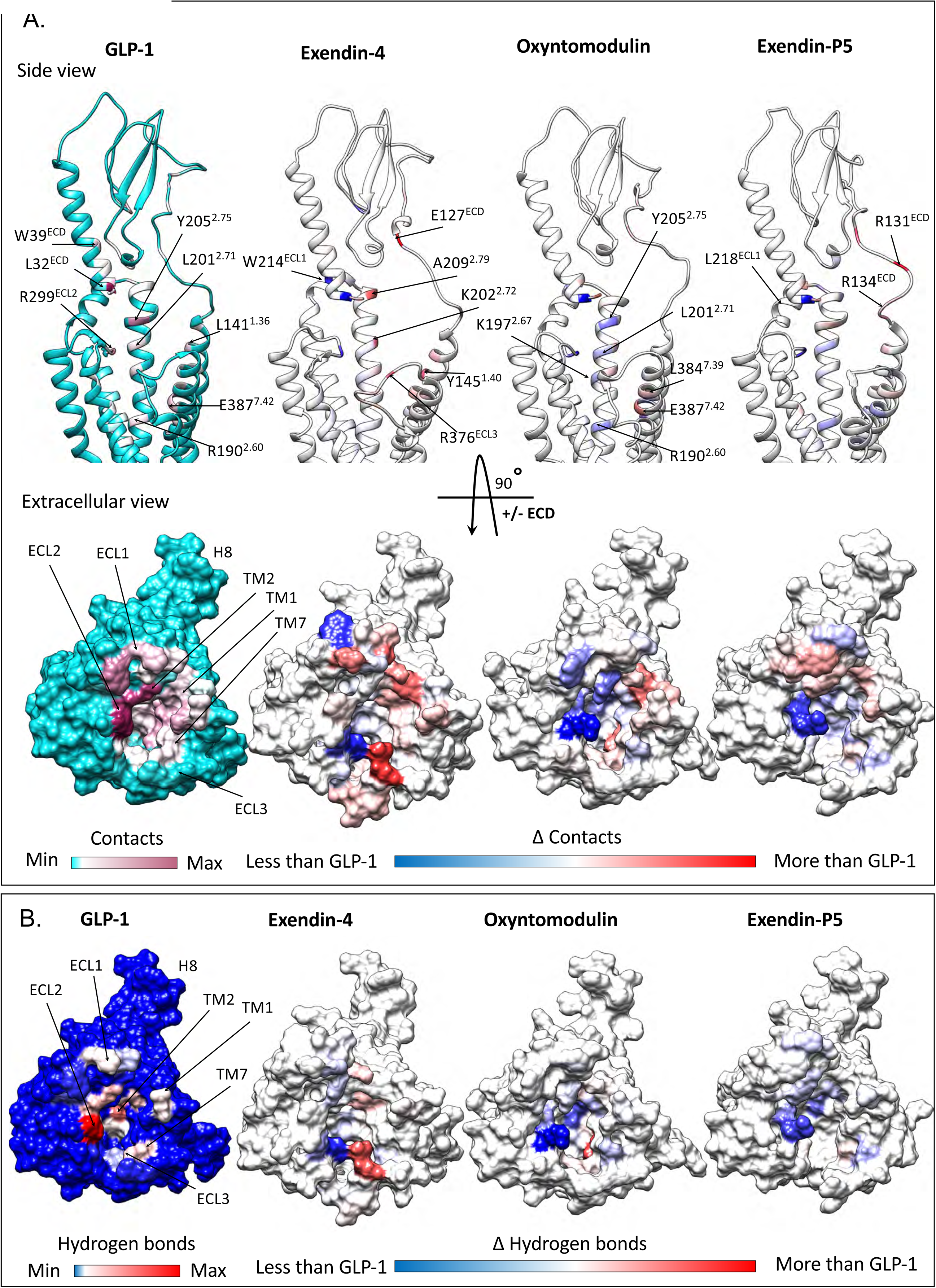
Contact differences in GLP-1R interactions of GLP-1, exendin-4, oxyntomodulin and exendin-P5 from MD simulations. **A,** Top; contact differences of each peptide with the GLP-1R TMD and ECD plotted on the receptor ribbon viewed from the side, Bottom; contact differences of each peptide with the TMD plotted on the GLP-1R surface viewed from the extracellular side (ECD not shown for clarity). The first column shows the GLP-1R contacts formed by GLP-1 during the simulations with no/min contacts in cyan and contacts heat-mapped from white to dark pink with increasing number/occupancy of interactions. The other three columns report the contacts differences for each residue of the GLP-1R during the MD simulation performed in the presence of other agonists with white indicating similar interactions to GLP-1, blue decreased contacts and red increased contacts. **B,** Hydrogen bond differences between the four peptides plotted on the surface of the TMD viewed from the extracellular side with the ECD removed for clarity. The first column shows the GLP-1R residues involved in hydrogen bonds with GLP-1 during the MD simulations with blue to red heatmap indicating the relative extent of interaction for each residue. The other three columns report the hydrogen bond differences for each GLP-1R residue during the MD performed in the presence of the other agonists with blue indicating less interactions, white similar and red more interactions, compared to GLP-1.

Consistent with the cryo-EM structure, the MD analysis revealed the GLP-1 N-terminus forms extensive interactions with TMs 1, 2, 3, 5, 6, 7 and ECLs 1-3 (Figure 6, Supplemental Figure 11). The interactions of oxyntomodulin differed, with more transient contacts deep within the peptide binding cavity and with residues along the peptide exposed face of TM2, ECL1 and ECL2. This was coupled with enhanced interactions at the top of TM1 and distinct and more sustained interactions within TM7 (Figure 6, Supplemental Table 3). Within the N-terminal nine amino acids, oxyntomodulin differs from GLP-1 by 2 residues, with Ala8 and Glu9 of GLP-1 replaced by Ser2 and Gln3 in oxyntomodulin (Figure 1A). These residues are located at the base of the GLP-1R binding pocket in the cryo-EM structures and interact with residues in TMs 1, 2 and 7 (Figures 2 & 3, Supplemental Figure 4, Supplemental Table 3). While Glu9 of GLP-1 formed strong and persistent hydrogen bond and van der Waals interactions with R190^2.60^, Y152^1.47^, and to a lesser extent, Y148^1.43^, the polar side chain of Gln3 in oxyntomodulin only formed weak and very transient hydrogen bonds with these residues, and van der Waals interactions only with Y148^1.43^ (Movie S1, Supplemental Table 3). Consistent with this, alanine substitution of Y148^1.43^ had a similar influence on the affinity of both ligands, while Y152^1.47^A had greater effect on GLP-1 affinity, and R190^2.60^A decreased GLP-1 affinity >30-fold, but oxyntomodulin was unaffected (Supplemental Figures 5 & 9, Supplemental Table 2). All three residues play a much greater role in GLP-1 mediated cAMP signalling, relative to oxyntomodulin (Figures 4-5, Supplemental Figures 7 & 10). In addition to these interactions, Gln3 of oxyntomodulin formed transient hydrogen bonds and hydrophobic contacts with T391^7.46^ and E387^7.42^, as well as van der Waals interactions with L388^7.43^, while Glu9 in GLP-1 only formed hydrophobic interactions with these residues. While Ser2 of oxyntomodulin and Ala8 of GLP-1 both interacted with TM7 residues, these interactions were stronger with oxyntomodulin due to a persistent hydrogen bond between Ser2 and E387^7.42^ and transient hydrogen bonds and van der Waals contacts with K383^7.38^, D372^ECL3^ and L384^7.39^ (Supplemental Table 3). Accordingly, alanine mutagenesis of TM7 residues influenced both GLP-1 and oxyntomodulin affinity, but there was a larger effect on oxyntomodulin, consistent with its stronger interactions, however, interestingly these residues were more important for GLP-1-mediated cAMP production (Supplemental Figures 9-10). In both peptides, the N-terminal histidine sits in an enclosed pocket forming hydrophobic and hydrogen bond interactions with E364^6.53^, E387^7.42^, R310^5.40^, Y241^3.44^, W306^5.36^, I313^5.43^ and Q234^3.37^ however, these interactions are more persistent with GLP-1, and this is likely linked to the stability of interactions of Glu9 and with residues at the base of the binding cavity (Supplemental Table 3, Supplemental Movie 1).

In the MD studies, weaker interactions of oxyntomodulin with residues deep in the cavity, particularly R190^2.60^, were coupled with a loss of stable interactions along the entire face of the TM2 helix and weaker interactions with ECL2, all of which interact with the same face of the helical GLP-1 peptide (Figure 6). While mutagenesis studies clearly reveal that these residues within TM2 and ECL2 are important for both GLP-1 and oxyntomodulin function, alanine mutations in these regions generally had a larger impact on GLP-1 (Figures 4-5, Supplemental Figures 9-10). Overall, these differing receptor interaction patterns of GLP-1 and oxyntomodulin were associated with larger conformational dynamics within the oxyntomodulin binding pocket during the course of the MD simulation, with the cavity opening and closing, likely due to the lack of stable interactions within the base of the TMD binding pocket, TM2 and ECL2, coupled with more persistent interactions with the upper regions of TM1 and TM7, relative to GLP-1 (Figure 6, Supplemental Figure 12, Supplemental Movie 1). While the C-terminus of oxyntomodulin and the receptor ECD/ECL1 was lower resolution in the cryo-EM map, the interaction patterns in the modelled protein and the MD simulations were largely similar to GLP-1, albeit that the nature of some interactions differed due to their differing sequences; residues capable of hydrogen bonding were present in one peptide, but not the other. For example, Arg17 and Arg18 of oxyntomodulin form interactions with the ECD and top of TM2 in the simulations, whereas the corresponding alanine residues in GLP-1 could not form these interactions. Interestingly the ECD was also more mobile in the oxyntomodulin bound receptor, when compared to GLP-1 (Supplemental Figure 12), which may be associated with the greater dynamics within the bundle, given the bound peptide bridges these two domains and stabilises their motions relative to one another.

The more open TMD in the static exendin-4 bound cryo-EM structure was associated with fewer stable peptide contacts (Supplemental Table 1), relative to GLP-1 and oxyntomodulin, however, the cryo-EM map was lower resolution, which precluded modelling of some of the binding site (top of TM6, ECL3 and ECL1). The MD simulations revealed a large number of conserved interactions between GLP-1 and exendin-4, including the majority of peptide hydrogen bonding within the TMD. Of particular note was the lack of persistent receptor interactions for His1 of exendin-4, in contrast to the extensive interactions observed for GLP-1 (His7) and oxyntomodulin (His1), as described above (Figure 3, Supplemental Tables 1 and 3, Supplemental Movie 1). His1 of exendin-4 formed only limited transient interactions with TM5, however more persistent interactions with E364^6.53^ in TM6 were evident in the MD simulations (Supplemental Table 3). Interestingly, His1-Gly2 of exendin-4 exhibited weaker density relative to the rest of the peptide in the cryo-EM map, and this is consistent with the enhanced flexibility observed for this region of the bound peptide in the simulations, likely due to Gly2, as glycine can destabilise α-helical conformations. The equivalent residue is a serine or alanine in oxyntomodulin and GLP-1 respectively, contributing to a more stable α-helical conformation, while also forming additional interactions at the base of the GLP-1R binding pocket (Figures 2-3, Supplemental Table 3). In line with strong and stable interactions of His1-Ala2 of GLP-1 compared with transient interactions of His1-Gly2 in exendin-4, truncation of these two N-terminal residues reduces GLP-1 affinity by 100-300 fold, whereas for exendin-4, this has no effect, albeit for both peptides, these residues are required for GLP-1R-mediated cAMP production^22–25^.

Despite the flexibility within the N-terminal two residues, Glu3 of exendin-4 formed very similar contacts to that of Glu9 of GLP-1, however, interactions with residues residing at the base of the pocket were more transient for exendin-4 (75-80% of the MD frames) than for GLP-1 in the MD simulations (97% of frames for Y152^1.47^ and 100% for R190^2.60^) (Supplemental Table 3). Receptor interactions for the remaining residues within the N-terminal 11 amino acids of exendin-4 were similar to those formed by GLP-1, albeit overall interactions deeper within TM2, TM3, TM5, and ECL2 were generally less persistent, and those in TM1 and TM7 were more persistent (Figure 6, Supplemental Table 3, Supplemental Movie 1). Similarity in their TMD interaction patterns is consistent with the strong correlation in the impact of TMD mutagenesis for these two agonists (Figure 5). Moreover, the more transient interactions of exendin-4 parallel the smaller effects of the mutagenesis on cAMP signalling for this peptide.

With the exception of His1, the largest difference in the interaction of GLP-1 and exendin-4 occurred within the mid region of the peptides where their sequences differ substantially (E^16^EAVRL^21^ for exendin-4 vs G^22^QAAKE^27^ for GLP-1). While the mid regions of both peptides interact with the top of TM1/ECD stalk, the ECD, ECL1 and the top of TM2, exendin-4 exhibits more persistent interactions in the simulations, particularly with the TM1/stalk and ECD (Supplemental Tables 1 and 3). This, in part, may account for the higher affinity of exendin-4 for the isolated receptor ECD and the higher affinity of amino terminally truncated exendin-4 peptides, where exendin(9-39) displays only 10-30-fold lower affinity compared to exendin-4, relative to the equivalent GLP-1(15-36), which has greater than 300-fold lower affinity than GLP-1^22, 31^. These interactions may also influence the conformation of TM1 accounting for the more outward conformation in the exendin-4 bound structure (Figure 3). Interestingly, single alanine amino acid substitutions of some interacting residues within the TMD had much larger effects on exendin-4 affinity than removal of the first 8 residues of the peptide, suggesting that peptide ECD and TMD interactions are correlated, such that non-optimal interactions with the TMD likely promote faster peptide dissociation from the ECD. This is also supported by previous studies, whereby Gly2Ala in exendin-4 was tolerated, but the converse for GLP-1 (Ala2Gly) reduced affinity^31–33^. However, the substitution of residues Glu22, Glu27 and Ala30 of GLP-1 with Gly16, Leu21 and Glu24 of exendin-4 enabled Gly2 tolerance^32^.

MD simulations on the exendin-P5-bound complex revealed a similar C-terminal interaction pattern to exendin-4 for this peptide, with more persistent interactions with the ECD and TM1/stalk than GLP-1 (Figure 6, Supplemental Figure 11, Supplemental Table 3). Nonetheless the TMD conformation differed and, overall, the receptor was relatively stable, exhibiting less flexibility compared with GLP-1R bound to exendin-4 or oxyntomodulin (Supplemental Figure 12). The N-terminal sequence of exendin-P5 differs considerably to the other peptides and is also extended by one residue (Figure 1A). Nonetheless, there are some commonalities with the other peptides in its pattern of interaction with the TMD. Val3-Asp4 interact with similar residues at the base of the binding cavity to those observed for residues 2 and 3 of the other peptides, however, with the exception of Y148^1.43^, these were very transient (Supplemental Table 3). Consequently, Asp4 could also form interactions with K197^2.67^ located higher within TM2. This provides a rationale for the effect of mutagenesis of R190^2.60^ and K197^2.67^, which were two of the few TMD residues that had a large impact on exendin-P5 affinity (Supplemental Figures 5 & 9). Despite occupying a different location in static structures, Glu1 of exendin-P5 interacts with multiple residues that interact with His1 of the other peptides, including E364^6.52^, E387^7.42^, R310^5.40^ and W306^5.36^. However, with the exception of R310^5.40^, where it forms a more persistent hydrogen bond, these interactions are again very transient and there are no interactions with TM3. Beyond these residues, there are very few interactions formed with the remainder of the N-terminal 9 residues of this peptide (Supplemental Table 3). Therefore, overall exendin-P5 forms relatively stable interactions with the ECD, upper portion of TM1, TM2 and TM7, and with ECL1 and ECL2, however interactions with key residues deeper in TM2, TM3, TM5, TM6, TM7, as well as with ECL3, are transient, the majority of them present for less than 20 % of the frames measured within the MD simulation (Supplemental Figure 11, Supplemental Table 3). These data are consistent with alanine substitution of TMD residues lining the binding cavity generally having limited impact on exendin-P5 affinity, suggesting it’s affinity is largely driven by interactions with the ECD (Figures 4-5, Supplemental Figure 9). In contrast, transient interactions between the exendin-P5 N-terminus and TMD are clearly important for agonism, with the majority of residues within this cavity being required for eliciting cAMP signalling (Figures 4-5, Supplemental Figure 10).

### Dynamics of peptide-TMD interactions are correlated with the allosteric effect of G proteins on agonist affinity and G protein conformation

The stark contrast in the requirement for stable TMD interactions for exendin-P5 affinity relative to GLP-1, oxyntomodulin and exendin-4, raises important questions regarding molecular mechanisms for peptide binding and receptor activation. To interrogate this, the influence of the bound G protein on the affinity of each agonist was assessed using wildtype and CRISPR-engineered HEK293 cells, where all Gα subtypes are depleted (*Δ*all G*α* HEK293)^34^. A NanoBRET membrane competition binding assay was employed to assess the ability of each peptide to inhibit binding of the fluorescent probe ROX-Ex4 to the GLP-1R N-terminally tagged with nanoluciferase (Nluc). In the wildtype cell line, GLP-1, oxyntomodulin and exendin-4 competition curves were clearly biphasic, with potencies for the high affinity site correlating with those reported from whole cell binding assays in the wildtype GLP-1R expressing ChoFlpIn cell line used in the mutagenesis study (Figure 7A, Supplemental Figures 5-6). In contrast, exendin-P5 exhibited monophasic binding curves with a lower pIC_50_ than the other peptides that was consistent with the pIC_50_ achieved in the ChoFlpIn whole cell assay. While there was a small reduction in the pIC_50_ for exendin-P5 in the *Δ*all G*α* cell line, this effect was relatively minor. In contrast, in the absence of G*α* proteins, the high affinity binding site for GLP-1, exendin-4 and oxyntomodulin was not observed; all displayed a single site for inhibition of Rox-Ex4 binding, and with IC_50_ values consistent with the lower affinity site in GLP-1R expressing membranes where G*α* proteins were present (Figure 7A). This loss of high affinity binding could be reversed by the overexpression of G_s_ in the *Δ*all G*α* cell line, consistent with G_s_ allosterically influencing the affinity of GLP-1, exendin-4 and oxyntomodulin, with much more limited effect on exendin-P5 (Figure 7A). Interestingly, when G_s_ was overexpressed, there was a larger influence on GLP-1 that exhibited highest stability of interactions within the TMD binding cavity in the MD studies, compared to oxyntomodulin and exendin-4, which were more dynamic (Figure 7A, Supplemental Movie 1).

**Figure 7.**
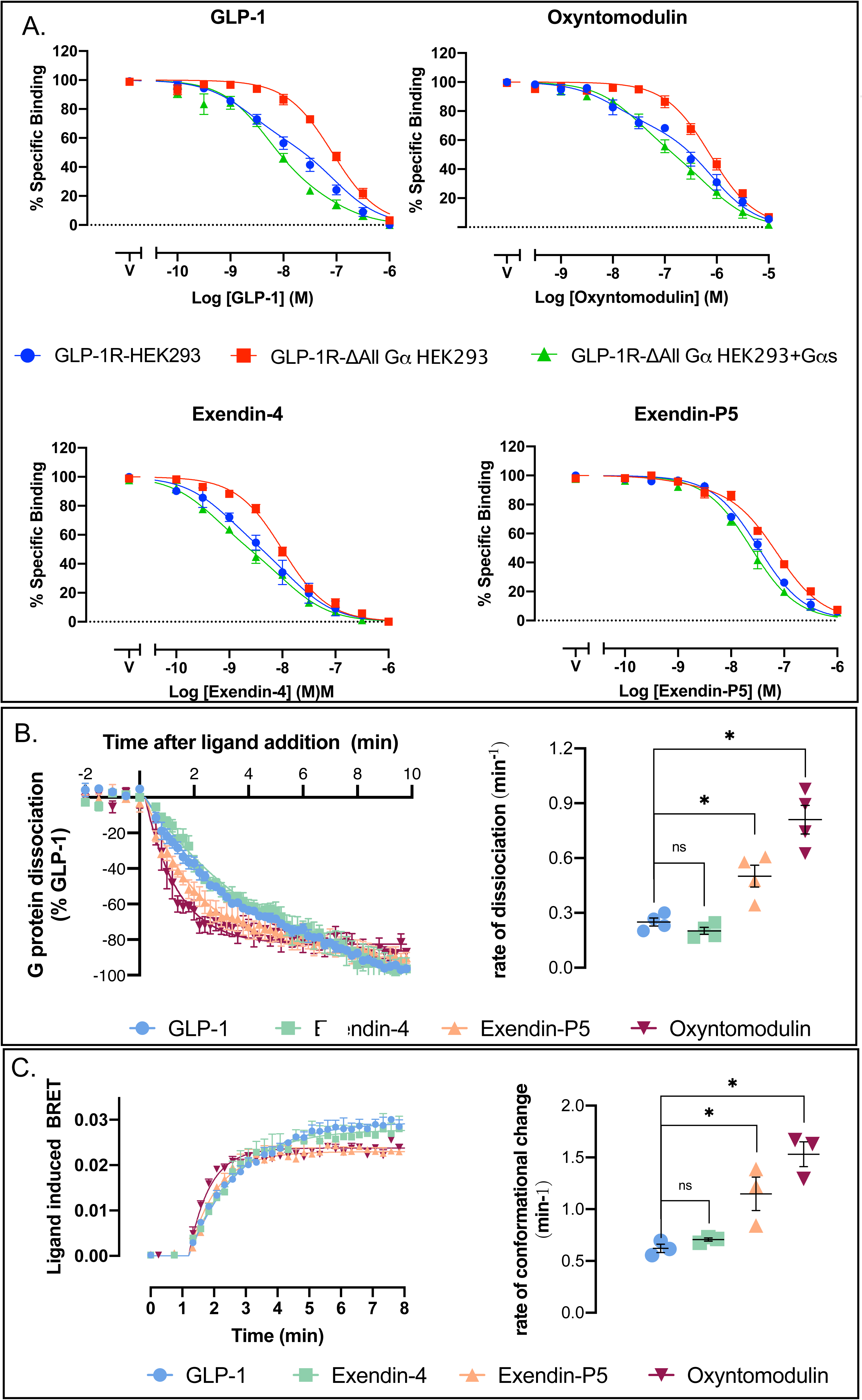
Allosteric effect of the G protein on peptide affinity and the peptide on G_s_ activation. **A,** Equilibrium competition binding assays assessing the ability of GLP-1, oxyntomodulin, exendin-4 and exendin-P5 to compete for the probe Rox-Ex4, in HEK293 cells overexpressing the GLP-1R in the presence of endogenous G*α* proteins (blue), the absence of G*α* proteins (red) and when G*α*s is overexpressed (no endogenous G*α* proteins) (green). Data are presented as % specific binding with 100% binding defined as total probe binding in the absence of competing ligand and non-specific (0%) binding determined as probe binding in the presence of 1μM exendin-4. Data are means ± s.e.m. of 5-7 independent experiments performed in duplicate. **B,** HEK293A cells transiently transfected with the GLP-1R and the NanoBit constructs for G*α*_s_ (G*α*-Lgbit, G*γ*_2_-Smbit). Left; Luminescence signal was assessed over time (0-20 min) in the presence of saturating concentrations of GLP-1 (1 μM), exendin-4 (1 μM), oxyntomodulin (10 μM) and exendin-P5 (10 μM) and responses were normalised to the max loss of luminescence observed with GLP-1. Data shown are mean ± s.e.m. of 4 independent experiments performed in triplicate. Right; Quantification of the rate of G protein dissociation (luminescence change) for each agonist was calculated by applying a one phase decay curve to the kinetic data with values from each individual experiment show in circles with the mean ± s.e.m. of the four individual experiments. **C,** Agonist induced changes in trimeric G_s_ protein conformation. Left; Ligand induced changes in BRET were measured in plasma membrane preparations performed in kinetic mode in the presence of saturating concentrations of GLP-1 (1μM), exendin-4 (1μM), oxyntomodulin (10μM) and exendin-P5 (10μM). Data shown are mean ± s.e.m. of 3 independent experiments performed in triplicate. Right; Quantification of the rate of ligand induced conformational change for each agonist was calculated by applying a one phase association curve to the kinetic data with values from each individual experiment shown in circles with the mean ± s.e.m. of the four individual experiments. * represents statistically different to GLP-1 (P<0.05) when assessed using a one-way ANOVA of variance with a Dunnett’s post hoc test.

A recent study revealed that while class A and class B GPCRs have similar kinetics for G protein recruitment, class B GPCRs exhibited slower G_s_ turnover associated with slower nucleotide exchange and slower GTP hydrolysis^21^. Collectively this manifested as slower ligand-induced dissociation of G*α* and G*βγ* when assessed in whole cells using a G_s_ Nanobit complementation assay. To assess if there is any potential for different GLP-1R agonists to display differences in G_s_ turnover, we employed this same Nanobit G_s_ complementation assay, to determine ligand-induced G protein dissociation in whole cells (Figure 7B). In addition, we also measured the kinetics of an earlier step in the G protein activation cycle using an assay to measure the G_s_ conformational change upon coupling to the ligand activated receptor (Figure 7C). This assay is sensitive to the positioning of the G*α*s *α*-helical domain (AHD), relative to the Ras homology domain (RHD) (performed in cell membranes in the absence of GTP), where separation of these domains is required for GDP release from the G protein^27, 35, 36^. Consistent with previous observations^13^, we demonstrated that exendin-P5 exhibited faster G_s_ conformational transitions, relative to GLP-1 and exendin-4, and this was coupled with faster dissociation of the G protein heterotrimer in the Nanobit assay (Figure 7B-C). Oxyntomodulin also displayed significantly faster kinetics relative to GLP-1 in both assays, whereas exendin-4 was more similar to GLP-1. In the G_s_ conformational assay, oxyntomodulin and exendin-P5 also exhibited a slightly lower maximal change in BRET than GLP-1 and exendin-4 (Figure 7C), suggesting a different ensemble of conformations of the AHD relative to the RHD, when bound by the different agonists.

### GLP-1R interactions with G_s_ are conserved, however MD simulations reveal ligand-specific effects on interaction dynamics

Overlay of the four consensus cryo-EM static structures revealed very similar backbone conformations of the intracellular face of receptor, with the greatest divergence for the ICLs (where modelled), and similar engagement with G_s_ in all four peptide bound structures (Figure 8). The C-terminal region of the *α*5 helix of G*α*_s_ was equivalently positioned for all structures, however, there was divergence at the N-terminal region of the *α*5, and this was translated across the remainder of the G protein (including the G*βγ* subunits). The MD simulations described above included the G_s_ heterotrimer and revealed that the receptor-G_s_ interactions were very similar regardless of the bound agonist, albeit the majority of interactions were transient (Figure 8B, Supplemental Figure 13, Supplemental Tables 4-5). Nonetheless, when exendin-4, oxyntomodulin and exendin-P5 were bound, the receptor exhibited less persistent hydrogen bonding with the G protein when compared to GLP-1. In addition, each complex displayed transient van der Waals interactions between the G*α* H4 and S6 domains and ICL3/TM6 of the receptor, that were, for the most part, not observed in the GLP-1 bound complex, albeit that specific interactions also differed between the individual peptide complexes (Figure 8B, Supplemental Figure 13, Supplemental Table 4). Moreover, while all the complexes displayed common and extensive interactions between the GLP-1R and the *α*5 helix of G*α*_s_, the persistence of interactions with individual residues differed in the exendin-P5 and oxyntomodulin bound complex (and to a lesser extent exendin-4), relative to GLP-1 over the time-course of the simulation (Figure 8B, Supplemental Table 4). Both the exendin-P5 and oxyntomodulin bound complexes also exhibited less persistent interactions between ICL2 and the *α*N and hns1/S1 region of G*α*_s_ and between the N-terminal portion of H8 with G*_β_*, whereas the remainder of interactions were largely consistent across the different complexes (Supplemental Figure 13, Supplemental Tables 4-5).

**Figure 8.**
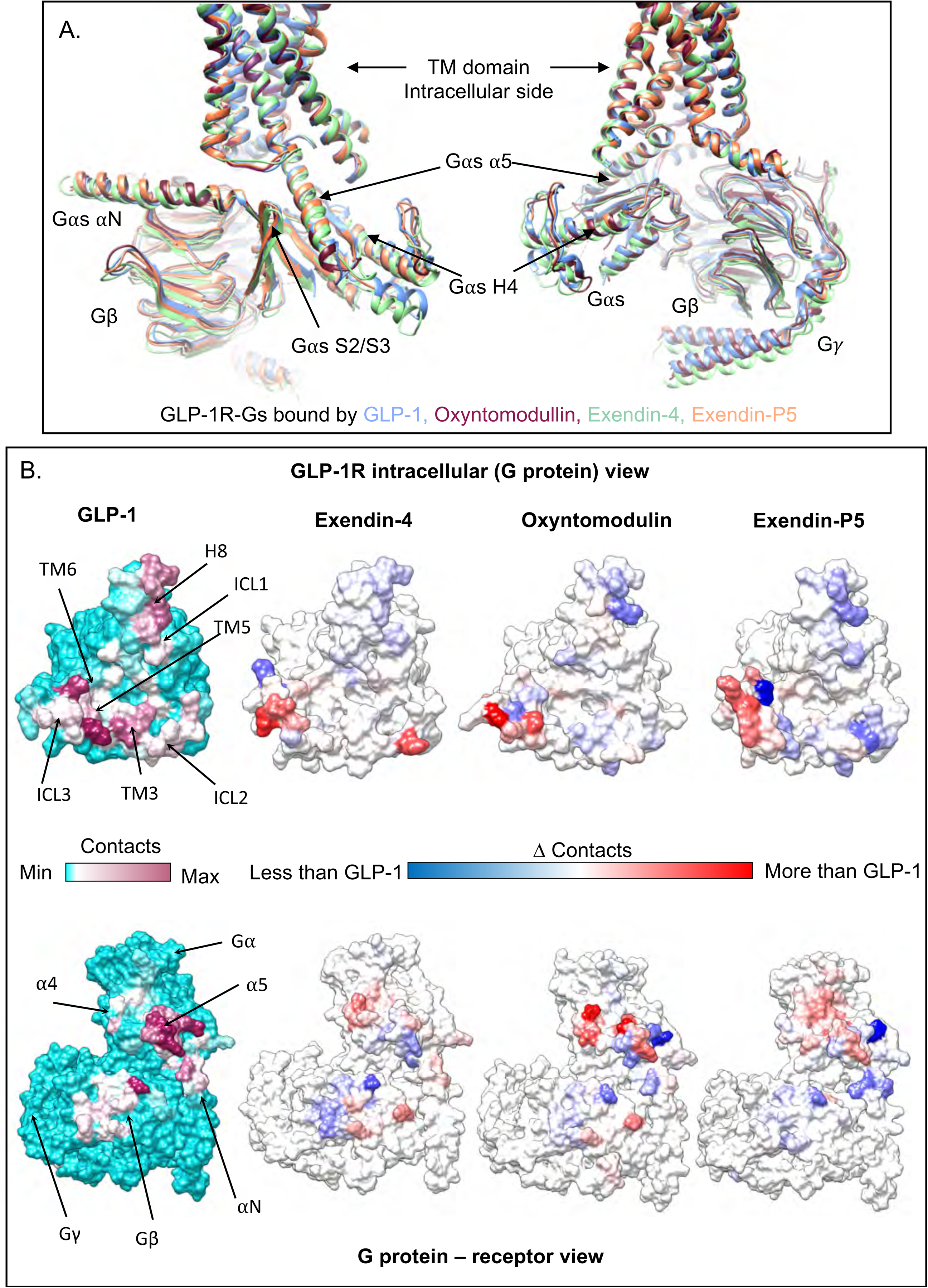
Interaction of G_s_ with the GLP-1R in the presence of different peptide agonists. **A,** Superimposition of GLP-1R:G_s_ structures bound with GLP-1 (PDB – 6X18, blue), exendin-4 (pale green), oxyntomodulin (dark pink) and exendin-P5 (pale orange), viewing the GLP-1R interface. **B,** Contact differences between the four complexes plotted on the receptor (top) and G_s_ (bottom) surface determined from MD simulations. The first column shows the contacts between GLP-1R (top) and G_s_ (bottom) during the MD simulations in the presence of GLP-1, with no contacts in cyan and increasing contacts heat mapped from white to dark pink. The other three columns report the contact differences (relative to GLP-1) for each residue of the GLP-1R and G_s_ during the MD performed in the presence of the other agonists with blue indicating less contacts, white similar contacts, and red enhanced contacts.

## Discussion

Combining experimentally determined GLP-1R structures with structure-function studies and simulations of receptor dynamics provides unique insights into how distinct agonists engage and activate the receptor. While GLP-1R peptide agonists bind both the ECD and the TMD, the role of the interactions with each domain differs among peptides. GLP-1, oxyntomodulin, and the clinically used mimetic exendin-4 are among the most extensively studied GLP-1R peptide agonists in functional and structure-function studies. Here, we reveal that while the peptides form similar interactions in the fully active, G_s_-coupled state of the receptor, exendin-4 and oxyntomodulin-occupied receptors are more dynamic, which may, in part, be linked to the more transient nature of their interactions with the TMD observed in MD simulations. Oxyntomodulin is a biased agonist relative to GLP-1^4^, and forms distinct and more dynamic interactions with the GLP-1R, particularly with residues at the base of the peptide binding cavity that are located above the conserved central polar network that is important for receptor activation. While the profile of exendin-4 mediated signalling is more similar to that of GLP-1, there are differences in receptor engagement by the two peptides that can be rationalised by the structural and dynamic data. Like oxyntomodulin, exendin-4 exhibits more transient interactions than GLP-1 with residues at the base of the peptide binding cavity, albeit it that the pattern of interactions with the polar core are relatively conserved.

G proteins can allosterically influence ligand affinity for GPCRs^37^. Consistent with this, we show the G protein can allosterically influence GLP-1R agonist affinity, but this occurs in a peptide-dependent manner. Enhanced affinity in the presence of G_s_ is correlated with the degree of closure of the extracellular side of the TMD cavity around the peptide N-terminus and also the dynamics of interactions with residues in this domain. GLP-1, oxyntomodulin and exendin-4, whose affinities are influenced by the presence of G_s_, promote a more closed bundle relative to exendin-P5, whose affinity is less sensitive to the presence of G_s_. N-terminally truncated GLP-1 and exendin-4 peptides lacking the first 8 amino acids (GLP-1(15-36) and exendin(9-39)), exhibit similar affinities in published studies^31^, to their corresponding full-length peptides in the absence of G protein, providing further support that the influence of G_s_ on peptide affinity is predominantly related to interactions of the N-terminus with the TMD. While it is likely that G_s_ has the potential to influence the affinity of all peptides that bind in the TMD cavity, the specific amino acid sequence of individual peptides impacts complementarity with receptor residues, influencing the stability of interactions and contributing to the degree of allosterically-facilitated TMD closure around the peptide. With the exception of interactions of Glu1 and Asp3 with polar residues deep in the bundle, the first 9 residues of exendin-P5 do not form stable interactions with the TMD, consequently the TMD is more open and interactions with the peptide-N-terminus play only a minor role in its overall affinity. Nonetheless, these transient interactions are clearly essential for receptor activation, with mutation of the majority of residues within the TMD cavity decreasing exendin-P5 efficacy, but not affinity.

Previously we revealed that the exendin-P5-bound GLP-1R induces faster G_s_ conformational transitions (that are linked to nucleotide exchange) and induces faster cAMP production when compared to the GLP-1-bound receptor^13^. In this study we demonstrate that both oxyntomodulin and exendin-P5 induce faster G_s_ conformational transitions and G_s_ heterotrimer dissociation, suggesting that these agonists may induce faster turnover of G_s_ than GLP-1 and exendin-4. Interestingly, this profile appears to be correlated to the strength and nature of interactions of these peptides with key polar residues at the base of the GLP-1R binding cavity. While GLP-1, and to a slightly lesser degree exendin-4, form very stable interactions with R190^2.60,^ Y152^1.47^, Y241^3.41^ and E364^6.53^, exendin-P5, and in particular, oxyntomodulin, form much more transient interactions with these key residues. This is predominantly due to the side chain chemistry of the residue at the position equivalent to Glu9 of GLP-1. While a glutamic acid is conserved in exendin-4, this is replaced by aspartic acid and glutamine in exendin-P5 and oxyntomodulin, respectively. As polar residues at these receptor locations are conserved across class B1 GPCRs and play a role in receptor activation for all receptors where studied^38–43^, stable vs transient interactions formed by peptide residues may also be associated with peptide efficacy at other Class B1 receptors.

Given cryo-EM structures of the GLP-1R in complex with G_s_ are stabilised by trapping the nucleotide-free G protein on the ligand-activated receptor, it is not surprising that the G_s_ interactions are similar across the four structures, even when bound by different agonists, nonetheless the MD simulations revealed potential differences in the dynamics of these interactions. Of particular note are the weaker interactions between ICL2 with the *α*N and hns1/S1 region of G*α*_s_ and between H8 and G*_β_*, in addition to differences in interactions between ICL3/TM6 within the *α*5 helix of the exendin-P5 and oxyntomodulin-bound complexes, compared to GLP-1. Interactions of G proteins with these domains contribute to separation of the G protein RHD and AHD, disruption of the P loop and the nucleotide binding pocket, all of which contribute to the release of GDP, one of the key rate-limiting steps in G protein activation^36^. However, how and if predicted differences in dynamics of receptor G protein interactions correlate to the differences in the effect of the allosteric coupling of the G protein on TMD binding sites, or the rates of G protein activation are unclear and additional studies will be required to address this.

Using the glucagon receptor as an exemplar, Hilger et al., identified that activated class B1 GPCRs exhibit a very persistent active state receptor conformation after G protein dissociation^21^. This implies that the activated receptor is primed to activate multiple rounds of G protein coupling following dissociation of the initial interacting protein, and this is proposed to contribute to the sustained cAMP signalling following activation of class B1 GPCRs. Consistent with this, exendin-P5, which induces faster kinetics in G protein activation and cAMP production, exhibits higher cAMP efficacy than GLP-1 and exendin-4, as indicated by a greater pEC_50_/pIC_50_ ratio when comparing cAMP and binding studies^13^. Our data suggests exendin-P5 may induce a “looser” coupling between the ligand TMD and G_s_ binding sites, potentially linked to the more short-lived interactions with TMD residues, resulting in higher efficacy due to faster G_s_ conformational transitions associated with nucleotide exchange and faster G_s_ dissociation. Overall, given the long-lived active receptor conformation, this would promote greater turnover of G_s_ and production of cAMP over time. A similar phenomenon has been observed at the calcitonin receptor, another class B1 GPCR, where human calcitonin, with a fast off-rate, turns over G protein faster than salmon calcitonin, which has a slow off rate, and this was related to differences in the residency time of the G protein on the receptor and to the sensitivity of the G protein to GTP^35^.

Oxyntomodulin exhibited even faster kinetics for G_s_ conformational transitions and subunit dissociation than those induced by exendin-P5, and this was also correlated with more transient interactions at the base of the binding cavity and TM2. However, in contrast, this ligand does not exhibit higher efficacy relative to GLP-1, as determined by comparing transduction ratios from cAMP signalling and their determined affinity measures (Supplemental Table 2). This highlights the complexity of GPCR activation, where downstream signalling is influenced by the interplay of multiple transducers that can interact with activated receptors (pleiotropic coupling), and can also be influenced by different trafficking profiles that alter the location of the receptor in the cell. Oxyntomodulin is a biased agonist relative to GLP-1^1, 3^ (and exendin-P5^26^), with bias towards arrestin recruitment over cAMP production, which would compete for G protein interactions and may contribute to the lack of correlation between the faster G_s_ dissociation and enhanced efficacy when assessing cAMP accumulation in whole cells over an extended timeframe. While the structural basis of GLP-1R biased agonism between arrestin recruitment and G protein pathways is still not clear and will require additional studies to decipher, there is growing evidence that biased agonism is associated with the conformation and dynamics of the TM6/ECL3/TM7/TM1 domain^3, 14, 16, 27, 29^, and this is consistent with the data presented herein. While in the consensus cryo-EM structures, GLP-1, exendin-4 and oxyntomodulin exhibit similar TMD conformations, MD simulations revealed that both exendin-4- and oxyntomodulin-bound GLP-1Rs, which exhibit biased agonism towards arrestin recruitment, are more dynamic in this region, than complexes with GLP-1 bound. In contrast, other agonists that exhibit stable open conformations, including exendin-P5, are correlated with bias towards G protein mediated signalling.

In summary, combining structural data from cryo-EM, receptor mutagenesis, pharmacological assays and MD simulations advances our understanding of peptide agonist engagement with the GLP-1R. As class B1 peptide agonists engage their receptors via a two-domain interaction, their efficacy for G protein mediated signalling is influenced by multiple factors, including the nature of interactions with the ECD and TMD, contributing to both peptide affinity and how ligand-receptor interactions influence G protein binding, nucleotide exchange and G protein dissociation from the receptor. This study provides insight into how differential dynamics of peptide-ligand engagement with the GLP-1R TMD can promote differences in G protein mediated signalling, improving molecular understanding of mechanisms that contribute to ligand-dependent differential efficacy at the GLP-1R.

## Methods

### Insect cell expression

HA-signal peptide-FLAG-3C-GLP-1R-3C-8*×*HIS^13^, human DNGα_s_^44^, His_6_-tagged human Gβ_1_and Gγ_2_ were expressed in *Tni* insect cells (Expression systems) using baculovirus as previously described. Cell cultures were grown in ESF 921 serum-free media (Expression Systems) to a density of 4 million cells/ml and then infected with three separate baculoviruses at a ratio of 2:2:1 for GLP-1R, DNGα_s_ and Gβ_1_γ_2_. Culture was harvested by centrifugation 60 h post infection and cell pellet was stored at -80°C.

### Complex purification

Cell pellet was thawed in 20 mM HEPES pH 7.4, 50 mM NaCl, 5 mM CaCls, 2 mM MgCl_2_ supplemented with cOmplete Protease Inhibitor Cocktail tablets (Roche) and benzonase (Merk Millipore). Complex formation was initiated by addition of 10 μM exendin-4 or 50 μM oxyntomodulin (China Peptides), Nb35–His (10 μg/mL) and apyrase (25 mU/mL, NEB); the suspension was incubated for 1 h at room temperature. The complex was solubilized from membrane by 0.5% (w/v) lauryl maltose neopentyl glycol (LMNG, Anatrace) supplemented with 0.03% (w/v) cholesteryl hemisuccinate (CHS, Anatrace) for 1 h at 4°C. Insoluble material was removed by centrifugation at 30,000 *g* for 30 min and the solubilised complex was immobilised by batch binding to M1 anti-FLAG affinity resin in the presence of 5 mM CaCl_2_. The resin was packed into a glass column and washed with 20 column volumes of 20 mM HEPES pH 7.4, 100 mM NaCl, 2 mM MgCl_2_, 5 mM CaCl_2_, 1 μM exendin-4 or 10 μM oxyntomodulin, 0.01% (w/v) LMNG and 0.0006% (w/v) CHS before bound material was eluted in buffer containing 5 mM EGTA and 0.1 mg/mL FLAG peptide. The complex was then concentrated using an Amicon Ultra Centrifugal Filter (MWCO 100 kDa) and subjected to size-exclusion chromatography on a Superdex 200 Increase 10/300 column (GE Healthcare) that was pre-equilibrated with 20 mM HEPES pH 7.4, 100 mM NaCl, 2 mM MgCl_2_, 1 μM exendin-4 or 10 μM oxyntomodulin, 0.01% (w/v) LMNG and 0.0006% (w/v) CHS to separate complex from contaminants. Eluted fractions consisting of receptor and G-protein complex were pooled and concentrated to 3-5 mg/mL. The complex samples were flash frozen in liquid nitrogen and stored at -80 °C.

### SDS–PAGE and Western blot analysis

Sample collected from size-exclusion chromatography was analysed by SDS–PAGE and Western blot. For SDS–PAGE, precast gradient TGX gels (Bio-Rad) were used. Gels were either stained by Instant Blue (Expedeon) or immediately transferred to PVDF membrane (Bio-Rad) at 100 V for 1 h. The proteins on the PVDF membrane were probed with two primary antibodies, rabbit anti-Gα_s_ C-18 antibody (cat. no. sc-383, Santa Cruz) against Gα_s_ subunit and mouse penta-His antibody (cat. no. 34660, QIAGEN) against His tags. The membrane was washed and incubated with secondary antibodies, 680RD goat anti-mouse and 800CW goat anti-rabbit (LI-COR). Bands were imaged using an infrared imaging system (LI-COR Odyssey Imaging System).

### Preparation of vitrified specimen

EM grids (Quantifoil, Großlöbichau, Germany, 200 mesh copper R1.2/1.3) were glow discharged for 30 s in high pressure air using Harrick plasma cleaner (Harrick, Ithaca, NY). Sample was applied on the grid in the Vitrobot chamber (FEI Vitrobot Mark IV). The chamber of Vitrobot was set to 100% humidity at 4°C. The sample was blotted for 5 s with a blot force of 20 and then plunged into propane-ethane mixture (37% ethane and 63% propane).

### Data acquisition

#### Exendin-4

Data for the GLP1:DNGs:exendin4 was collected on a Titan Krios microscope operated at 300 kV (ThermoFisher Scientific equipped with a Gatan Quantum energy filter operating in Zero Loss mode with a energy slit with of 20 eV and a Gatan K2 summit direct electron detector (Gatan). Movies were taken in EFTEM nanoprobe mode, with 50 µm C2 aperture and no objective aperture, at a magnified pixel size of 0.87 Å. Each movie comprised 48 frames with a total dose of 48 e-/Å2, exposure time was 8 s with a dose rate of 7 e-/pix/s on the detector. EPU (ThermoFisher Scientific) was used to automate data collection which involved implementing beam-tilt to collect a 3×3 grid of holes. As the data collection was split between two different days, the data was split into 18 optics groups.

#### Oxyntomodulin

Data was collected on a Titan Krios microscope operated at 300 kV (ThermoFisher Scientific equipped with a Gatan Quantum energy filter and a Gatan K2 summit direct electron detector (Gatan) and a Volta Phase Plate (ThermoFisher Scientific). Movies were taken in EFTEM nanoprobe mode, with 50 µm C2 aperture, at a magnified pixel size of 1.06 Å. Each movie comprised 50 frames with a total dose of 50 e-/Å2, exposure time was 8 s with a dose rate of 7 e-/pix/s on the detector. Data acquisition was done using SerialEM software at -500 nm defocus^44^.

### Data processing

#### Exendin-4

8816 movies were collected and subjected to motion correction using motioncor2^45^. CTF estimation was done using Gctf software^46^ on the non-dose-weighted micrographs. The particles were picked from dose-weighted and low-pass filtered micrographs using crYOLO automated picking routine^47^. The particles were extracted in RELION 3.0^48^ using a box size of 256 pixels. 1.52 M picked particles were subjected to rounds of 2D and 3D classification in order to obtain a homogenous set of projections. This led to 422 k particles which were polished and had their CTF parameters re-refined in RELION. Further rounds of 2D and 3D classification yielded a final particle stack of 277.5 k particles for final 3D refinement and further rounds of masked refinements to reveal details of more flexible regions of the protein. Final consensus refinement produced a structure resolved to 3.59 Å (FSC = 0.143, gold standard). The cryo-EM data collection, refinement and validation statistics are reported in Supplemental Data Table 6.

#### Oxyntomodulin

Data processing of oxyntomodulin: 2,364 movies were collected and subjected to motion correction using motioncor2^45^. Contrast transfer function (CTF) estimation was done using Gctf software on the non-dose-weighted micrographs^46^. The particles were picked using gautomatch (developed by K. Zhang, MRC Laboratory of Molecular Biology, Cambridge, UK; http://www.mrc-lmb.cam.ac.uk/kzhang/Gautomatch/). An initial model was made using EMAN2^49^ based on a few automatically picked micrographs and using the common-line approach. The particles were extracted in RELION 2.03^50^ using a box size of 180 pixels. Picked particles (1070980) were subjected to two rounds of 3D classification with 3 classes. Particles (209000) from the best looking class were subjected to 3D auto-refinement in RELION 2.03. The refined revealed the final structure at 3.3 Å resolution. The cryo-EM data collection, refinement and validation statistics are reported in Supplemental Data Table 6.

### Modelling

The sequence corrected model of exendinP5-GLP-1R-Gs (PDB: 6B3J)^13^ was used as the initial template and fit in the cryo-EM density maps in UCSF Chimera (v1.14) for both the exendin-4 and oxyntomodulin bound structures, followed by molecular dynamics flexible fitting (MDFF) simulation with nanoscale molecular dynamics (NAMD)^51^. The fitted models were further refined by rounds of manual model building in COOT^52^ and real space refinement, as implemented in the PHENIX software package^53^. The ECD and ECLs were modelled manually without ambiguity based on the ECD-focused map. Density of ECD linker (E127^ECD^-P137 ^ECD^), ECL1 (T207^ECL1^-Q213^ECL1^/L218^ECL1^) and ICL3 (N338^ICL3^-D344^ICL3^) regions of both exendin 4 and oxyntomodulin complexes were discontinuous and these sequences were omitted from the final models. The ECL3 was less resolved in exendin-4 complex and residues from E373^ECL3^ to R376^ECL3^ were omitted.

### ChoFlpIn stable cell lines generation

The wild-type (WT) and mutant cMyc-GLP-1R^4^ constructs containing designed signal alanine mutation were integrated into CHOFlpIn cells using the FlpIn Gateway technology system (Invitrogen). Stable CHOFlpIn expression cell lines were selected using 600 μg/ml hygromyocin B, and maintained in DMEM supplemented with 5% (V/V) FBS (Invitrogen) at 37°C in 5% CO_2_.

### Whole cell radioligand binding assay

Cells were seeded at a density of 30,000 cells/well into 96-well culture plates and incubated overnight in DMEM containing 5% FBS at 37°C in 5% CO_2_. Growth media was replaced with binding buffer [DMEM containing 25 mM HEPES and 0.1% (w/v) BSA] containing 0.1 nM [^125^I]-exendin(9–39) and increasing concentrations of unlabelled peptide agonists. Cells were incubated overnight at 4°C, followed by three washes in ice cold 1*×* PBS to remove unbound radioligand. Cells were then solubilised in 0.1 M NaOH, and radioactivity determined by gamma counting. For all experiments, nonspecific binding was defined by 1 μM exendin(9– 39).

### cAMP accumulation assay

CHOFlpIn WT GLP-1R or CHOFlpIn mutant GLP-1R cells were seeded at a density of 30,000 cells per well into a 96-well plate and incubated overnight at 37 °C in 5% CO_2_. cAMP detection was using a Lance cAMP kit (PerkinElmer Life and Analytical Sciences), performed as previously described^6^. Growth media was replaced with stimulation buffer [phenol-free DMEM containing 0.1% (w/v) bovine serum albumin (BSA) and 1 mM 3-isobutyl-1-methylxanthine] and incubated for 1 h at 37 °C in 5% CO_2_. Cells were stimulated with increasing concentrations of ligand, 100 μM forskolin or vehicle, and incubated for 30 min at 37°C in 5% CO_2_. The reaction was terminated by rapid removal of the ligand-containing buffer and addition of 50 μL of ice-cold 100% ethanol. After ethanol evaporation, 75 μL of lysis buffer [0.1% (w/v) BSA, 0.3% (v/v) Tween 20, and 5 mM HEPES, pH 7.4] was added, and 10 μL of lysate was transferred to a 384-well OptiPlate (PerkinElmer Life and Analytical Sciences). 5 μL of 1/100 dilution of the Alexa Fluor® 647-anti cAMP antibody solution in the Detection Buffer (PerkinElmer Life and Analytical Sciences) and 10 μL of 1/5000 dilution of Eu-W8044 labeled streptavidin and Biotin-cAMP in the Detection Buffer (PerkinElmer Life and Analytical Sciences) were added in reduced lighting conditions. Plates were incubated at room temperature for 2 h before measurement of the fluorescence using an EnVision Multimode Plate (Reader PerkinElmer Life and Analytical Sciences). All values were converted to cAMP concentration using cAMP standard curve performed parallel and data were subsequently normalized to the response of 100 μM forskolin in each cell line, and then normalized to the WT for each agonist.

### Cell surface expression

Cell surface expression was detected using a cell surface ELISA to detect a double c-Myc epitope label incorporated with the N-terminal region of the GLP-1R constructs using the protocol described previously^54^.

### NanoBRET membrane binding assays

HEK293A WT and HEK293A ΔGα cells were transiently transfected with Nluc-hGLP-1R or Nluc-hGLP-1R + Gα_s_ + G*β*1*γ*2. 48 h post transfection, cells were harvested and plasma membrane was extracted as described previously^35^. 2 μg per well of cell membrane was incubated with furimazine (1:1,000 dilution from stock) in assay buffer (1× HBSS, 10 mM HEPES, 0.1% (w/v) BSA, 1× P8340 protease inhibitor cocktail, 1 mM DTT and 0.1 mM PMSF, pH 7.4). Rox-Ex4 was used as the fluorescent ligand in the NanoBRET binding assay. Membranes were increasing concentrations of peptides and RoxEx4 for 30 minutes prior to measurement of the BRET signal between Nluc-hGLP-1R and Rox-Ex4. This was assessed using a PHERAstar (BMG LabTech) at 10 sec intervals (25 °C). A Kd concentration, (3.16 nM for HEK293 WT cells, and 10 nM for HEK293 ΔGα cells) of Rox-Ex4 was used. Data were corrected for baseline and vehicle (probe only) responses.

### G_s_ conformational change

HEK293A cells stably expressing the GLP-1R (tested and confirmed to be free from mycoplasma) were transfected with a 1:1:1 ratio of Gγ_2_:venus–Gα_s_:nanoluc–Gβ_1_ using a standard PEI protocol (6:1 ratio PEI:DNA). Cells were incubated overnight at 37°C in 5% CO_2_. Collection and preparation of cell plasma membranes and assessment of G protein conformational change was performed using a previously established method^27, 35^. Briefly, 5 µg per well of cell membrane was incubated with furimazine (1:1,000 dilution from stock) in assay buffer (1× HBSS, 10 mM HEPES, 0.1% (w/v) BSA, 1× P8340 protease inhibitor cocktail, 1 mM DTT and 0.1 mM PMSF, pH 7.4). The GLP-1R-induced BRET signal between Gα_s_ and Gγ was measured at 30°C using a PHERAstar (BMG LabTech). Baseline BRET measurements were taken for 2 min before addition of vehicle or ligand. BRET was measured at 15 s intervals for a further 10 min. All assays were performed in a final volume of 100 μl. Data were corrected for baseline and vehicle control. The concentration response curves were then plotted using the total area under the curve during the time of measurement post ligand addition.

### G_s_ Nanobit complementation assays

HEK293AWT cells stably expressing the hGLP-1R were transiently transfected with G*α*-LgBIT, G*β*1, G*γ*2-SmBIT (1:5:5) 48 h before the assays using standard PEI transfection protocol. Cells were then incubated with coelenterazine H (5 μM) in assay buffer (1× HBSS, 10 mM HEPES, 0.1% (w/v) BSA) for 1 h at room temperature. Luminance signals were measured using a CLariostar (BMG LabTech) at every 30 s intervals before, and every 15 s intervals after ligand addition (25°C). Data were corrected to baseline and vehicle treated samples.

### Pharmacological data analysis

Pharmacological data were analysed using Prism 8 (GraphPad). Concentration response signalling data were analysed as previously described using a three-parameter logistic equation. cAMP accumulation concentration-response curves were analysed using an operational model of agonism modified to directly estimate the ratio of *τ*/*K_A_* as described previously^55^.

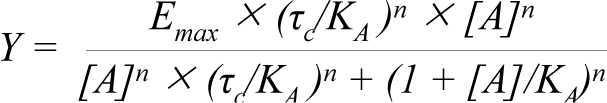

where *E_m_* represents the maximal stimulation of the system, *K_A_* is the agonist-receptor dissociation constant, in molar concentration, [*A*] is the molar concentration of ligand and *τ* is the operational measure of efficacy in the system, which incorporates signalling efficacy and receptor density. Derived *τ*/*K_A_* values were corrected to cell surface expression, measured by ELISA, and errors were propagated from both *τ*/*K_A_* and cell surface expression.

For rate analysis of G protein BRET assays, data were fitted to a one-phase association curve. Normalised AUC for the indicated ligand concentrations were plotted as a concentration response curve and fitted with a three-parameter logistic curve.

Statistical analysis was performed using a one-way analysis of variance and a Dunnett’s post-test, and significance accepted at P < 0.05.

### MD methods

The missing loops in the cryo-EM structures were reconstructed using modeller or by molecular superposition as described elsewhere^27^. The four GLP-1R complexes were prepared for MD simulations with the CHARMM36 force field^56^, employing in-house python htmd^57^ and TCL (Tool Command Language) scripts. Hydrogen atoms were first added at a simulated pH of 7.0 by means of the pdb2pqr^58^ and propka^59^ software, and the protonation of titratable side chains was checked by visual inspection. Each system was superimposed on the GLP-1R coordinates retrieved from the OPM database^60^ in order to correctly orient the receptor before it was inserted^61^ in a rectangular 125 Å x 116 Å 1-palmitoyl-2-oleyl-sn-glycerol-3-phosphocholine (POPC) bilayer (previously built by using the VMD Membrane Builder plugin 1.1, Membrane Plugin, Version 1.1. at http://www.ks.uiuc.edu/Research/vmd/plugins/membrane/), removing the lipid molecules overlapping the receptor TMs bundle. TIP3P water molecules^62^ were added to the simulation box (125 Å x 116 Å x 195 Å) by means of the VMD Solvate plugin 1.5 (Solvate Plugin, Version 1.5. at <http://www.ks.uiuc.edu/Research/vmd/plugins/solvate/). Overall charge neutrality was finally reached by adding Na^+^/Cl^-^ counter ions (final ionic concentration of 0.150 M), using the VMD Autoionize plugin 1.3 (Autoionize Plugin, Version 1.3. at <http://www.ks.uiuc.edu/Research/vmd/plugins/autoionize/).

### Systems equilibration and MD settings

Equilibration and MD productive simulations were computed using ACEMD. Isothermal-isobaric conditions (Berendsen barostat^63^ with a target pressure 1 atm; Langevin thermostat^64^ with a target temperature 300 K and damping of 1 ps^-1^) were employed to equilibrates the systems through a multi-stage procedure (integration time step of 2 fs). First, clashes between lipid atoms were reduced through 3000 conjugate-gradient minimization steps, then a 2 ns long MD simulation was run with a positional constraint of 1 kcal mol^-1^ Å^-2^ on protein and lipid phosphorus atoms. Successively, 20 ns of MD were performed constraining only the protein atoms. In the last equilibration stage, positional constraints were applied only to the protein backbone alpha carbons, for a further 60 ns.

Four 500 ns-long replicas were simulated for each complex (2 μs of total MD time). Productive trajectories were computed with an integration time step of 4 fs in the canonical ensemble (NVT) at 300 K, using a thermostat damping of 0.1 ps^-1^ and the M-SHAKE algorithm^65^ to constrain the bond lengths involving hydrogen atoms. The cutoff distance for electrostatic interactions was set at 9 Å, with a switching function applied beyond 7.5 Å. Long range Coulomb interactions were handled using the particle mesh Ewald summation method (PME)^66^ by setting the mesh spacing to 1.0 Å.

### MD Analysis

Atomic contacts were computed using VMD^13^. A contact was considered productive if the distance between two atoms was lower than 3.5 Å. Hydrogen bonds were quantified using GetContacts analysis tool (at https://getcontacts.github.io/), a donor-acceptor distance of 3.3 Å and an angle value of 150° were set as geometrical cut-offs. The video was produced employing VMD^67^ and avconv (at https://libav.org/avconv.html).

### Graphics

Molecular graphics images were produced using the UCSF Chimera (v1.14) and ChimeraX packages from the Computer Graphics Laboratory, University of California, San Francisco (supported by NIH P41 RR-01081and R01-GM129325)^68, 69^.

## Data availability statement

All relevant data are available from the authors and/or are included with the manuscript or Supplemental Information. Atomic coordinates and the cryo-EM density map have been deposited in the Protein Data Bank (PDB) under accession numbers 7LLY and 7LLL, and EMDB entry IDs EMD-23436 and EMD-23425, for oxyntomodulin and exendin-4, respectively.

## Acknowledgements

This work was supported by the National Health and Medical Research Council of Australia (NHMRC) (project grant 1126857, ideas grant 1184726 and SRF 1160076 (D.W.)), and program grant 1150083 (P.M.S.)). P.M.S. is a Senior Principal Research Fellow and D.W. a Senior Research Fellow of the NHMRC. R.D. was supported by Takeda Science Foundation 2019 Medical Research Grant and Japan Science and Technology Agency PRESTO (18069571). K.J.G., K.L and P.Z. are Future Fellows of the Australian Research Council (K.J.G - FT170100392, K.L - FT160100075, P.Z – FT200100218). CAR is a Royal Society Industry Fellow. This work was supported the Monash University Ramaciotti Centre for cryo-electron microscopy, and the Monash University MASSIVE high-performance computing facility.

## Author contributions

G.D designed, performed and analysed the MD simulations; Y-L.L and X.Z expressed and purified the complexes; M.K and R.D vitrified the samples; M.K and H.V imaged the samples to acquire EM data; X.Z and M.K processed the EM data and performed EM map calculations; X.Z, M.J.B and A.K. built the models and performed refinements; L.C and T.T.T performed the mutagenesis studies; P.Z and T.T.T performed the G protein and binding studies; G.D, X.Z, M.J.B, P.Z, P.M.S and D.W performed data analysis and data interpretation; Y-L.L, A.G, K.J.G, K.L, A.C, R.D and C.A.R assisted with data interpretation, and supervision for the project; P.Z, P.M.S and D.W designed and supervised the project, and wrote the manuscript with assistance from G.D, X.Z and M.J.B. All authors reviewed and revised the manuscript.

## Competing interests

The authors declare no competing interests.

---

**Supplemental Figure 1.**
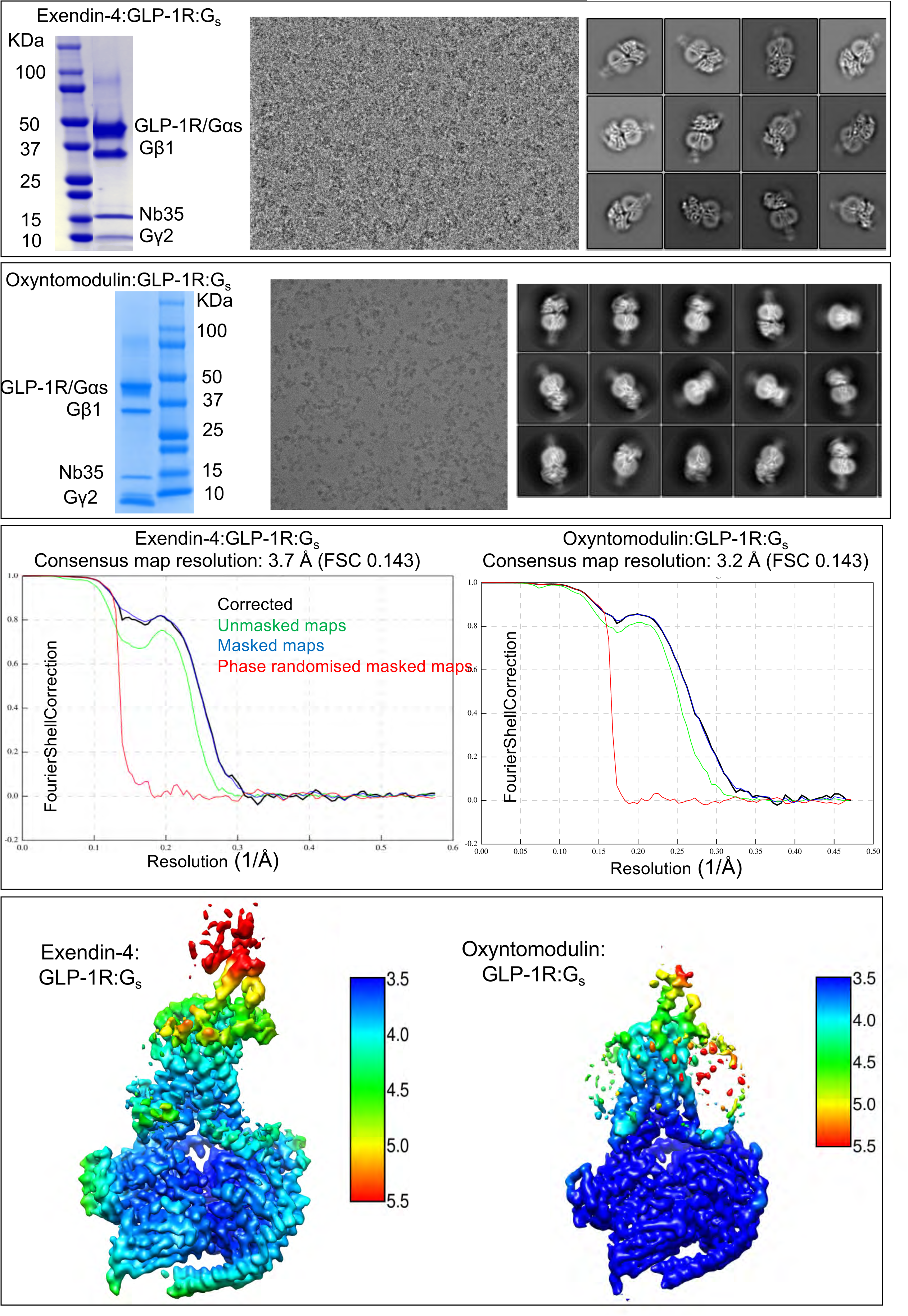
Purification, cryo-EM data imaging and processing of exendin-4 and oxyntomodulin bound GLP-1R:G_s_ complexes. **A,** Exendin-4 bound complex. Left; Coomassie stained gel showing the prep purity containing all expected components. Middle; Representative micrograph of the exendin-4:GLP-1R:G_s_ complex. Right; Two-dimensional class averages of the complex in LMNG micelle. **B,** oxyntomodulin bound complex. Left; Coomassie stained gel showing the prep purity containing all expected components. Middle; Representative micrograph of the oxyntomodulin:GLP-1R:G_s_ complex. Right; Two-dimensional class averages of the complex in LMNG micelle. **C,** “Gold standard” Fourier shell correlation (FSC) curves for the exendin-4 bound complex (left) and the oxyntomodulin bound complex (right), showing the overall nominal resolution at 3.7 Å and 3.3 Å, respectively. **D,** Cryo-EM density maps for the exendin-4 complex (left) and the oxyntomodulin bound complex (right), coloured according to local resolution (scaled in Å; dark blue, highest resolution, red, lowest resolution).

**Supplemental Figure 2.**
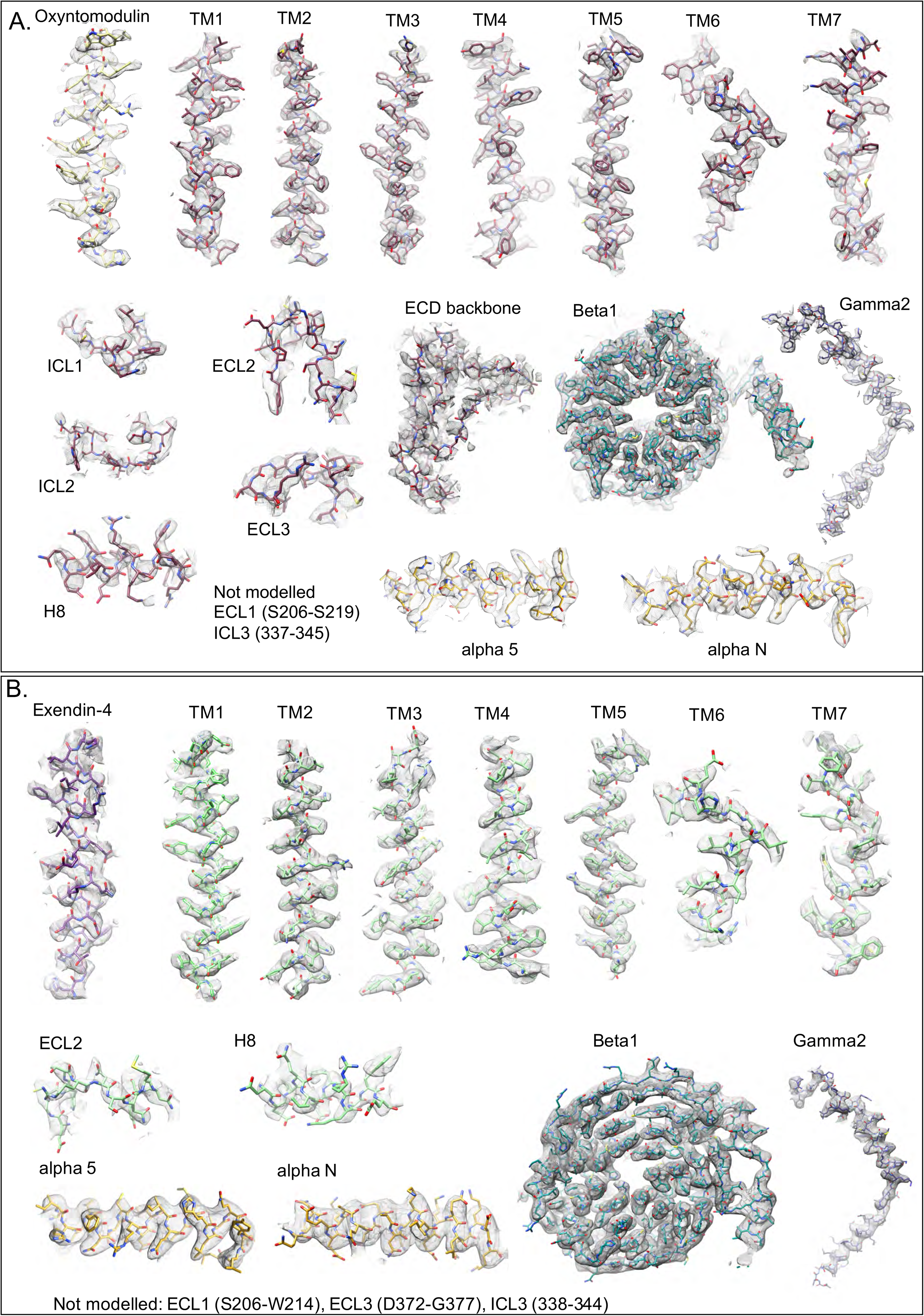
The atomic resolution model of the exendin-4-bound and oxyntomodulin bound GLP-1R:Gs in the cryo-EM density map. EM density map and the model are shown for all seven TM helices and H8 of the receptor, ECLs and ICLs where modelled, the α5 and α5 helices of the Gα_S_ Ras-like domain, *β*1 and *γ*2 and the peptide agonists for the oxyntomodulin (top) and exendin-4 (bottom) bound complexes. The ECD density to model fit is also shown for the oxyntomodulin sample.

**Supplemental Figure 3.**
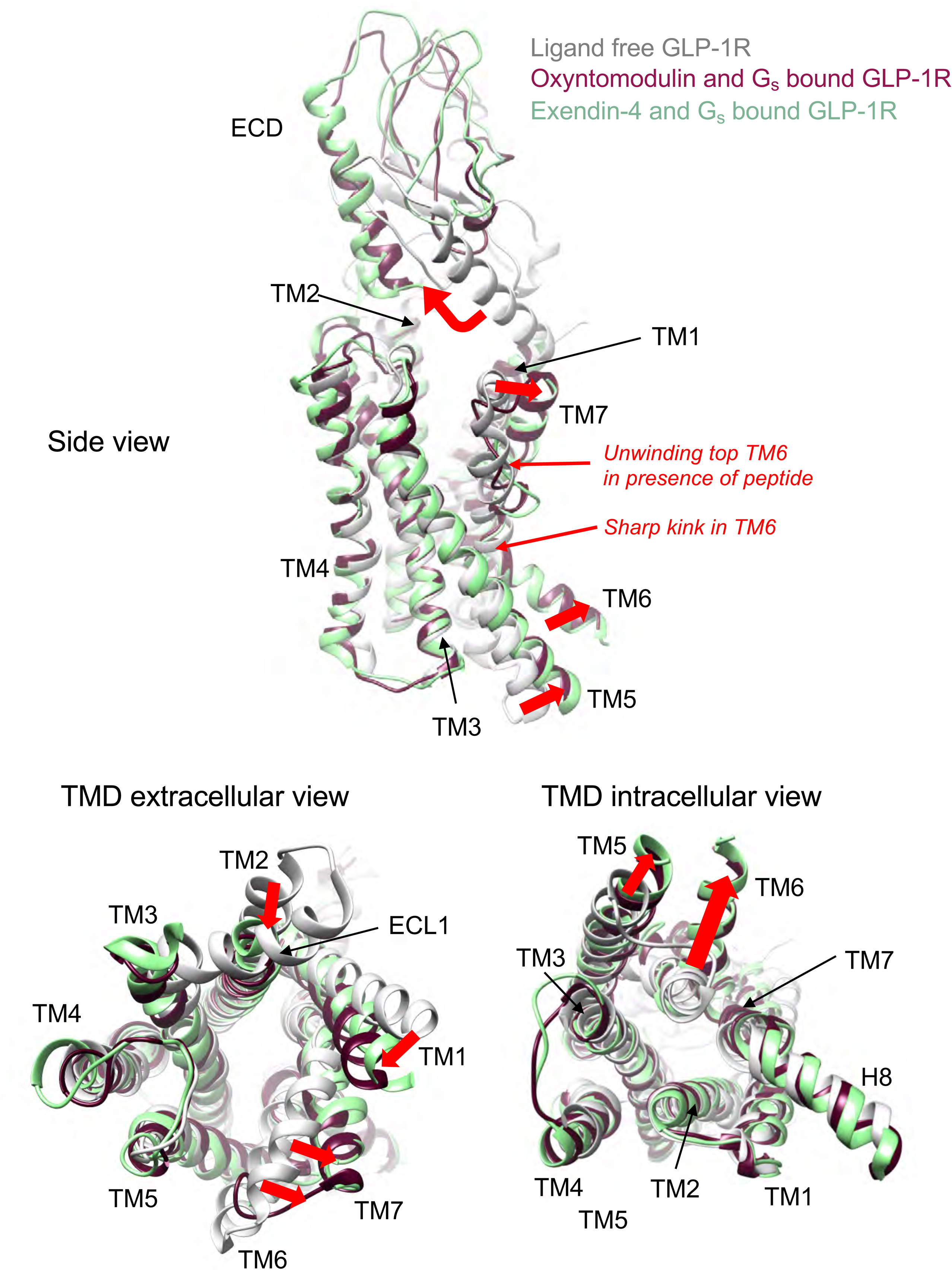
GLP-1R conformational transitions from the inactive conformation to G_s_ bound conformations induced by exendin-4 and oxyntomodulin. Superimposition of the inactive ligand free GLP-1R structure (6LN2^30^ – pale grey) with the GLP-1R in its G_s_ bound conformation induced by oxyntomodulin (dark pink) and exendin-4 (pale green). Top; full length receptors viewed from the side. Bottom left; TMD viewed from the extracellular face. Bottom right; TMD viewed from the intracellular face. Red arrows/labels highlight the most substantial backbone conformational transitions between the inactive (grey) and the activated peptide bound (dark pink/pale green) GLP-1R conformations.

**Supplemental Figure 4.**
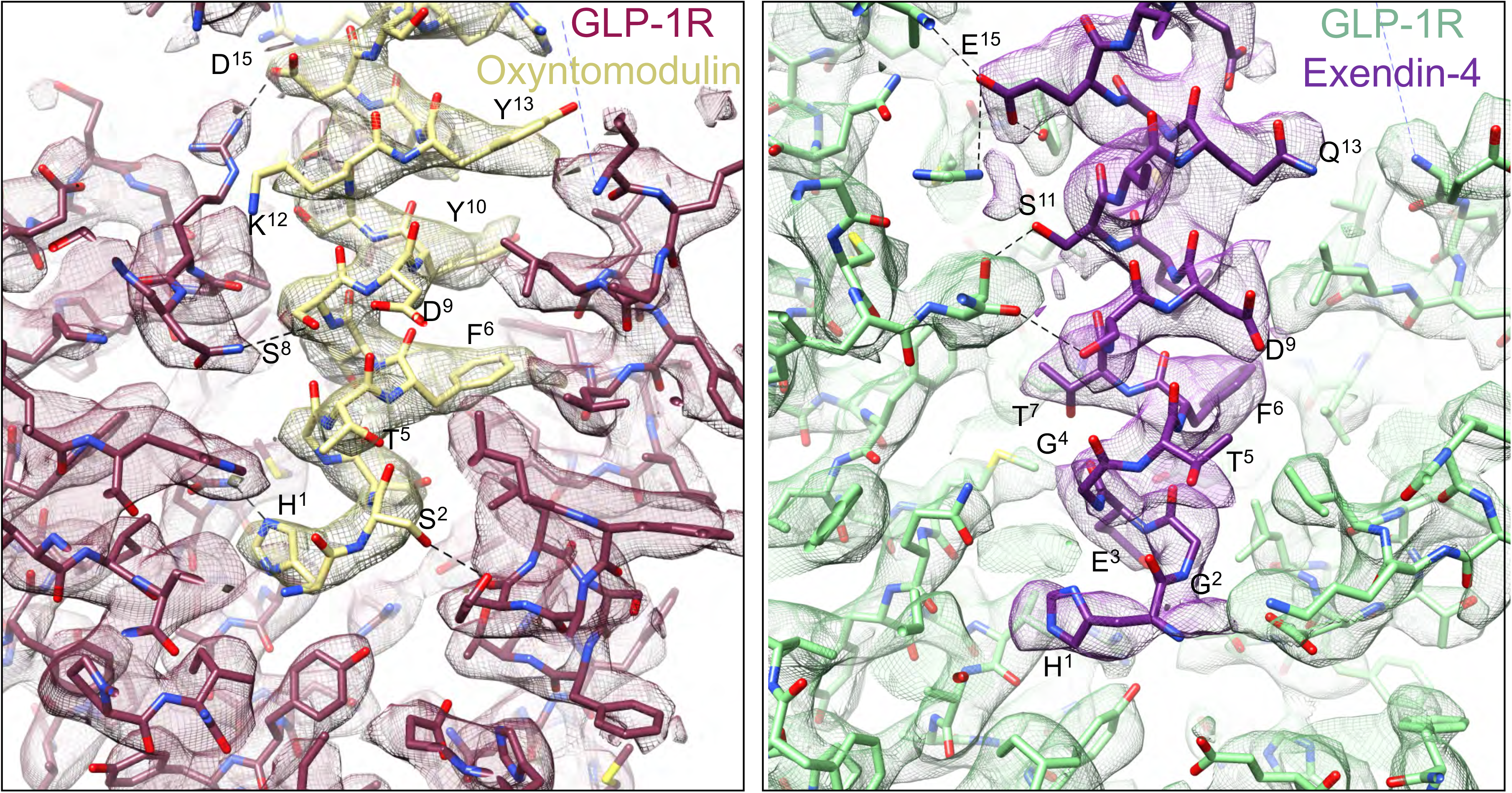
Cut through of the peptide binding cavity in the TMD showing the models and cryo-EM maps. Cut-through of the TMD peptide binding cavity highlighting the density on the peptide and the interacting residues for the oxyntomodulin bound GLP-1R (left) and the exendin-4 bound GLP-1R (right). Residues in the peptide are labelled. *Linked to Figure 2* where peptide and interacting TMD residues are labelled.

**Supplemental Figure 5.**
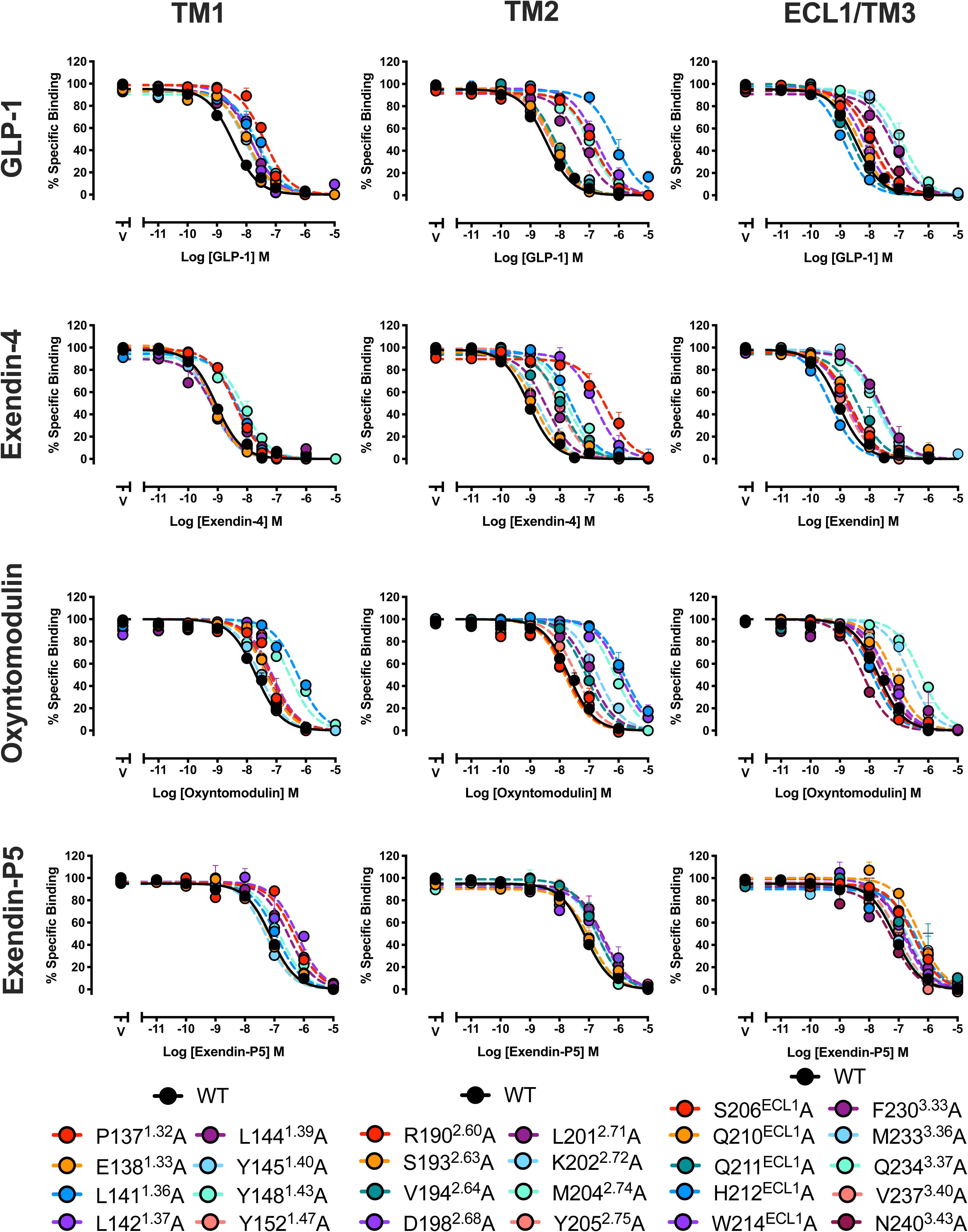
Inhibition binding curves for wildtype and alanine mutants of GLP-1R residues within TMs 1-3. Equilibrium competition binding assays assessing the ability of GLP-1, oxyntomodulin, exendin-4 and exendin-P5 to displace the radioligand ^125^I-exendin(9-39), in Cho-FlpIn cells overexpressing wildtype or mutant GLP-1Rs. Data are presented as % specific binding with 100% binding defined as total probe binding in the absence of competing ligand and non-specific (0%) binding determined as probe binding in the presence of 10 μM exendin(9-39). Data are means + s.e.m. of 4-10 independent experiments for mutant receptors (wildtype = 11-35 individual experiments) performed in duplicate.

**Supplemental Figure 6.**
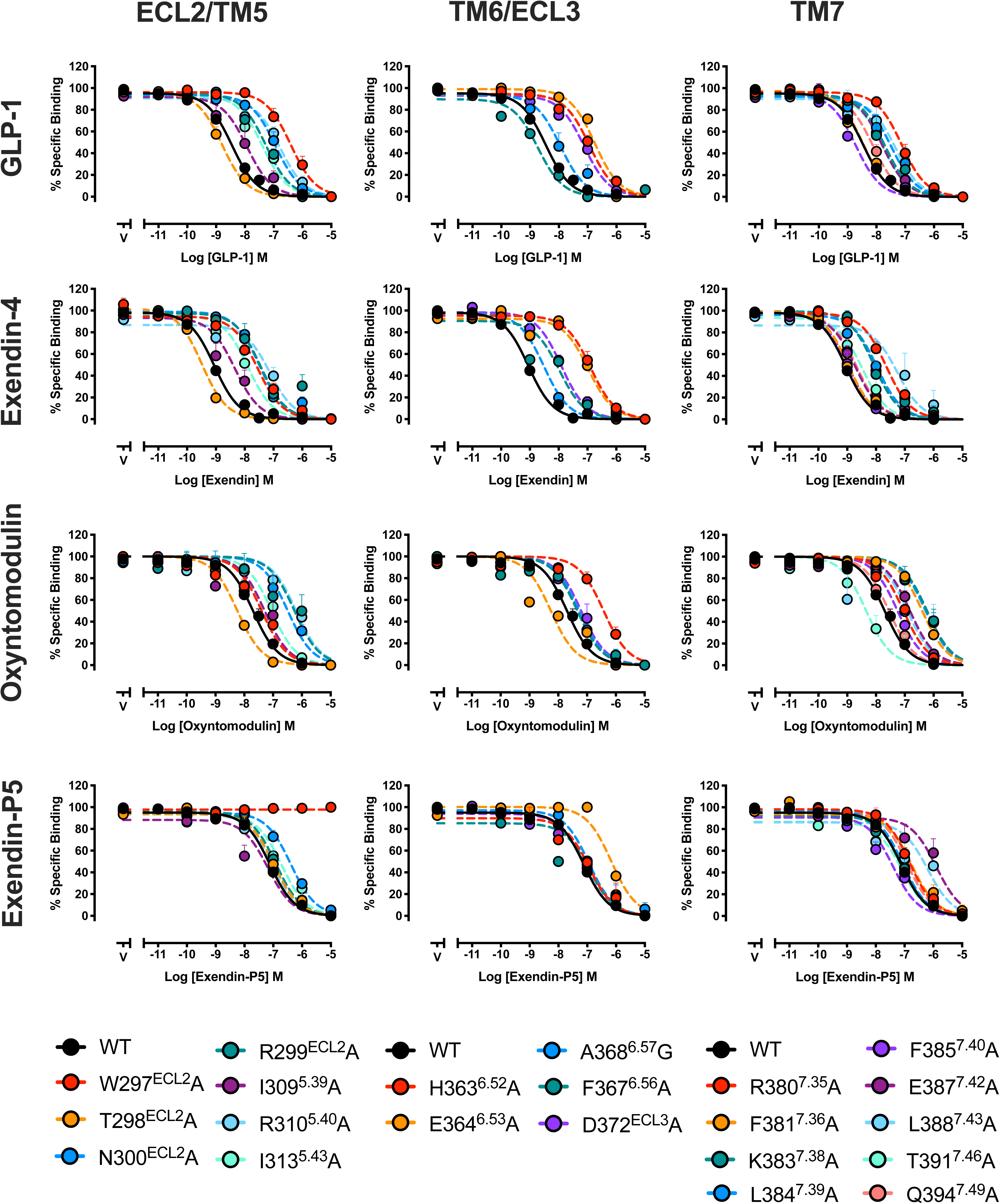
Inhibition binding curves for wildtype and alanine mutants of GLP-1R residues within TMs 4-7. Equilibrium competition binding assays assessing the ability of GLP-1, oxyntomodulin, exendin-4 and exendin-P5 to displace the radioligand ^125^I-exendin(9-39), in Cho-FlpIn cells overexpressing wildtype or mutant GLP-1Rs. Data are presented as % specific binding with 100% binding defined as total probe binding in the absence of competing ligand and non-specific (0%) binding determined as probe binding in the presence of 10 μM exendin(9-39). Data are means + s.e.m. of 4-10 independent experiments for mutant receptors (wildtype = 11-35 individual experiments) performed in duplicate.

**Supplemental Figure 7.**
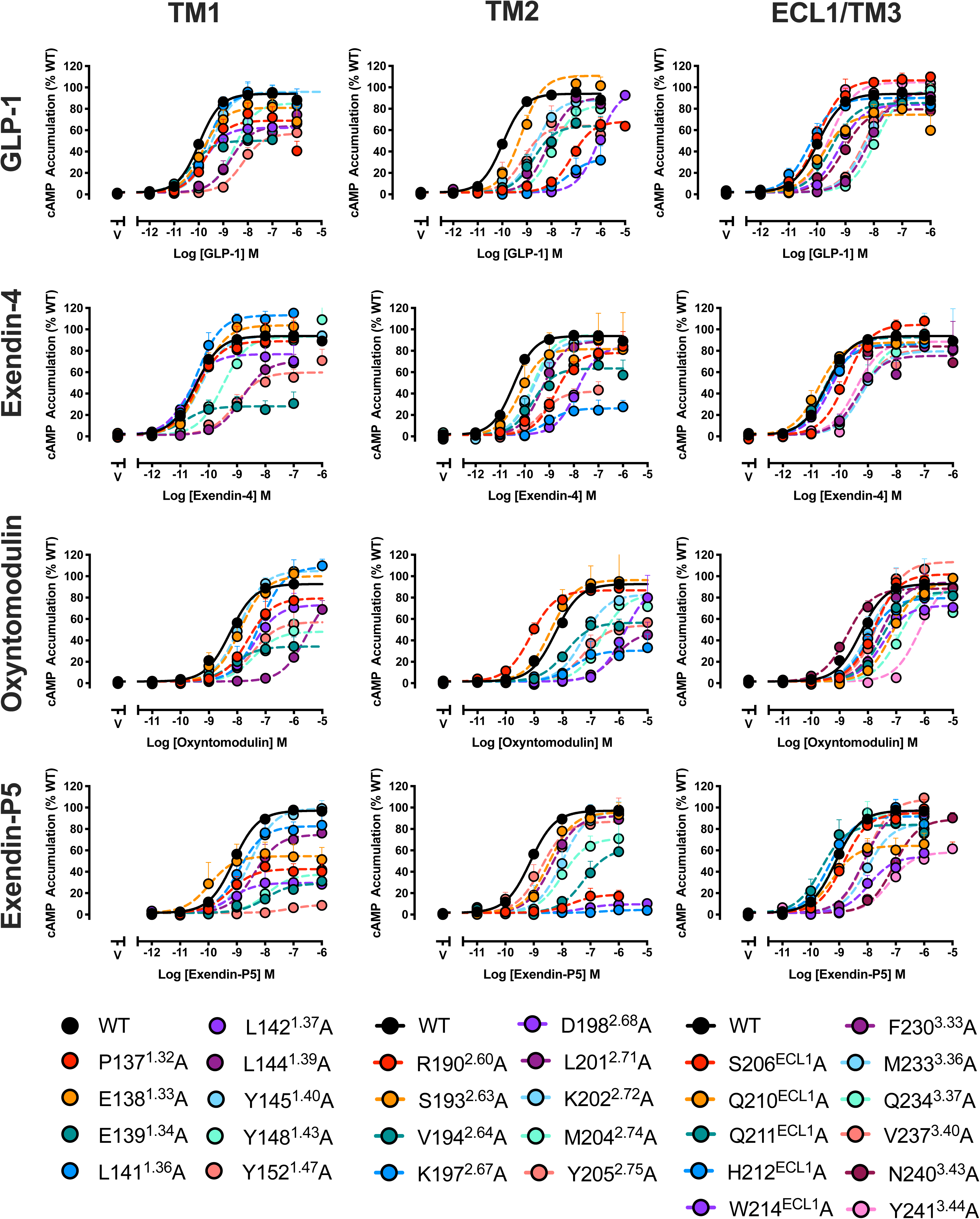
cAMP accumulation in ChoFlpIn cells expressing wildtype or alanine mutants of GLP-1R residues within TMs 1-3. Concentration response curves for cAMP production by GLP-1, oxyntomodulin, exendin-4 and exendin-P5 in Cho-FlpIn cells overexpressing wildtype or mutant GLP-1Rs. Data are presented as % cAMP accumulation mediated by the wildtype receptor. Data are means + s.e.m. of 5-11 independent experiments for mutant receptors (wildtype = 36-45 individual experiments) performed in duplicate.

**Supplemental Figure 8.**
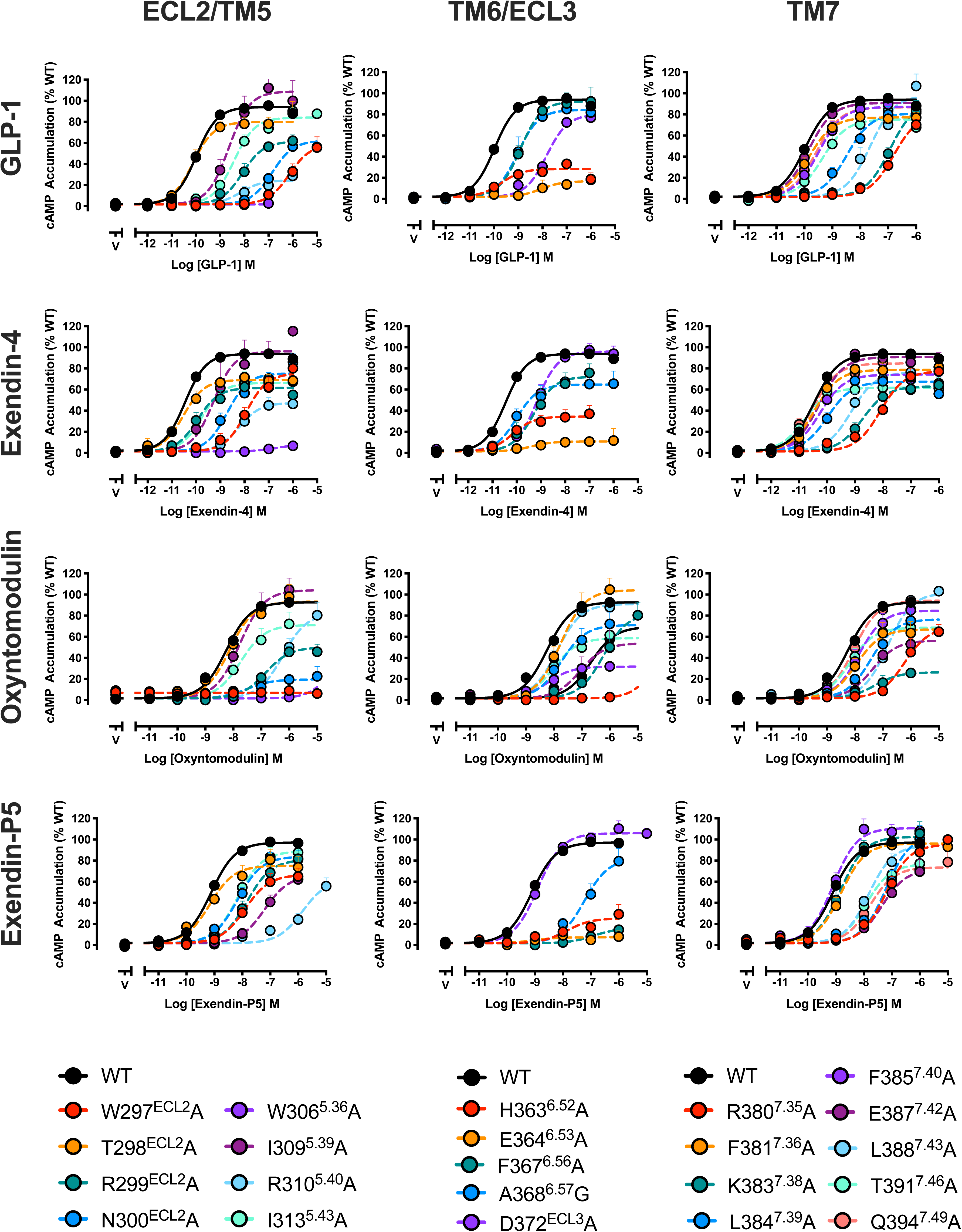
cAMP accumulation in ChoFlpIn cells expressing wildtype or alanine mutants of GLP-1R residues within TMs 4-7. Concentration response curves for cAMP production by GLP-1, oxyntomodulin, exendin-4 and exendin-P5 in Cho-FlpIn cells overexpressing wildtype or mutant GLP-1Rs. Data are presented as % cAMP accumulation mediated by the wildtype receptor. Data are means + s.e.m. of 5-11 independent experiments for mutant receptors (wildtype = 36-45 individual experiments) performed in duplicate.

**Supplemental Figure 9.**
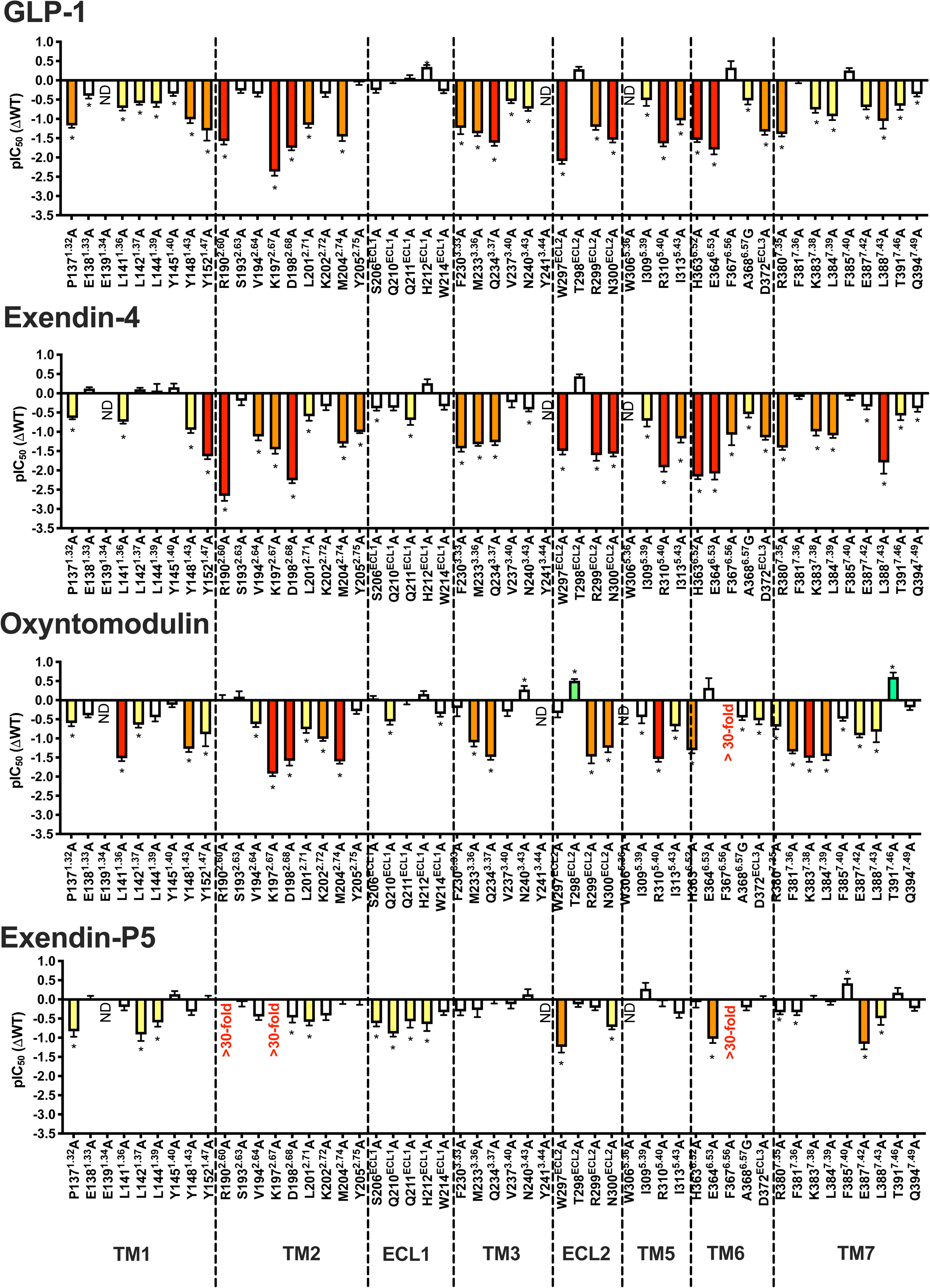
Agonist affinity profiles of GLP-1R alanine mutants reveal the importance of individual residues for peptide affinity. pIC_50_ values for each peptide were derived from radioligand inhibition binding experiments from data in Supplemental Figures 5-6. Bars represent differences in calculated affinity (pIC_50_) values for each mutant relative to the wild-type receptor for GLP-1, exendin-4, oxyntomodulin and exendin-P5. Statistical significance of changes in affinity in comparison with wild-type was determined by one-way analysis of variance and Dunnett’s post-test, and are indicated with an *asterisk* (*, *p* < 0.05). Data that are statistically significant are coloured based on the extent of effect. All values are mean + s.e.m. of 4-10 independent experiments, conducted in duplicate. ND, not determined. *Also see Figures 4-5*.

**Supplemental Figure 10.**
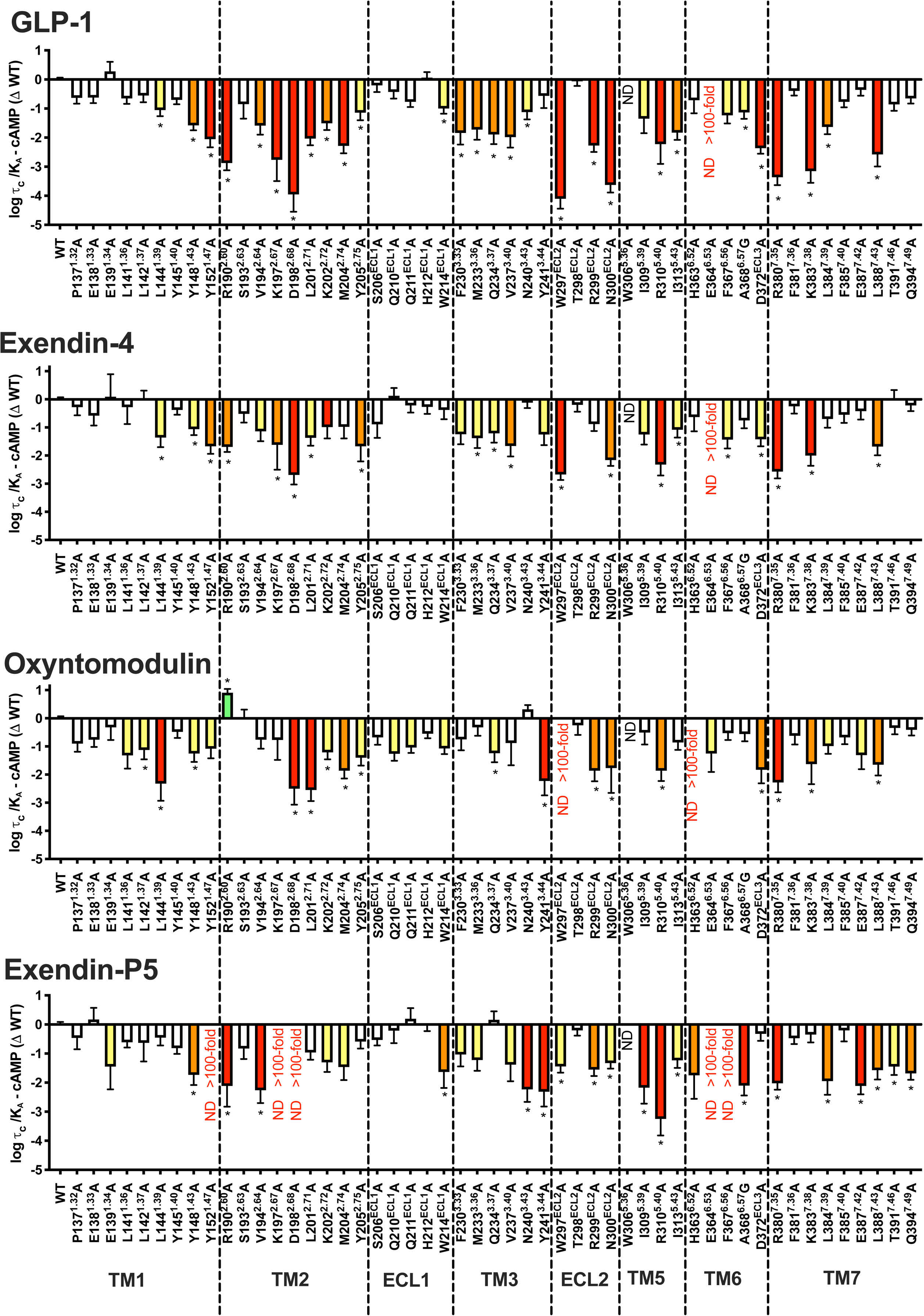
Peptide-dependent effects of TMD alanine mutations on cAMP signalling. Differences in the coupling efficiency (log(τ*/K_A_)_c_*) for cAMP formation of TMD mutations compared to the wild-type receptor by GLP-1, exendin-4, oxyntomodulin and exendin-P5 were determined by applying the operational model of agonism to concentration response data shown in supplemental figures 7-8. Statistical significance of changes in coupling efficacy was determined by one-way analysis of variance and Dunnett’s post-test, and those of significance are indicated with an *asterisk* (*, *p* < 0.05 compared with wild-type). Data that are statistically significant are coloured based on the direction and extent of effect. All values are mean + s.e.m. of 5-11 independent experiments, conducted in duplicate. ND. Not determined.

**Supplemental Figure 11.**
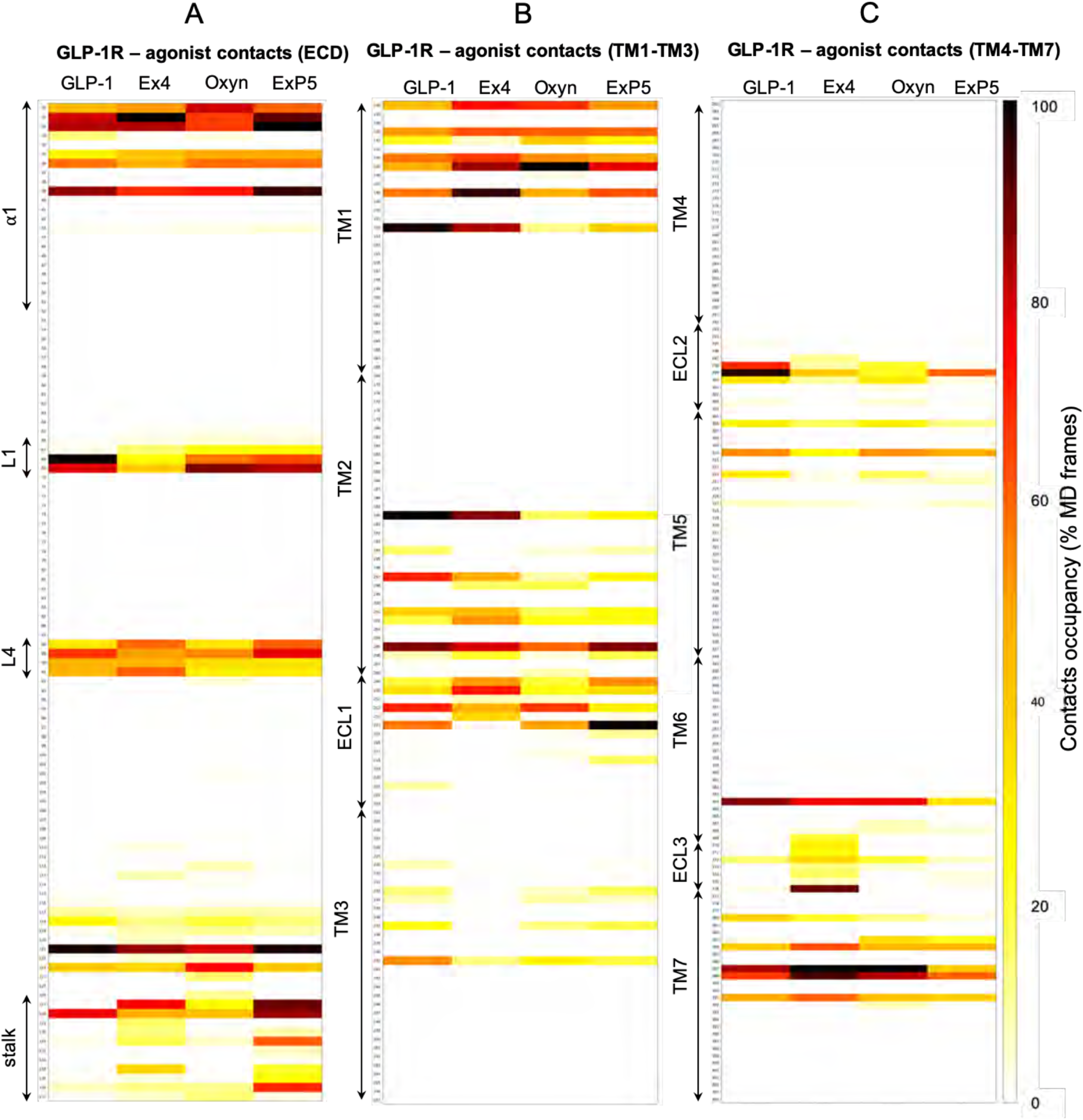
Heatmap showing the GLP-1R contacts and peptide agonists from MD simulations. The GLP-1R primary sequence is divided into three segments: ECD (A), TM1 to TM3 (B), and TM4 to TM7 (C). ICL1, ICL2, and IC3 have been omitted as they are not involved in interactions. For each agonist (GLP-1, exendin-4 (Ex4), oxyntomodulin (Oxyn), exendin-P5 (ExP5)), data are reported as the percentage of MD frames (occupancy) with at least one interatomic contact and normalized for the maximum occupancy.

**Supplemental Figure 12.**
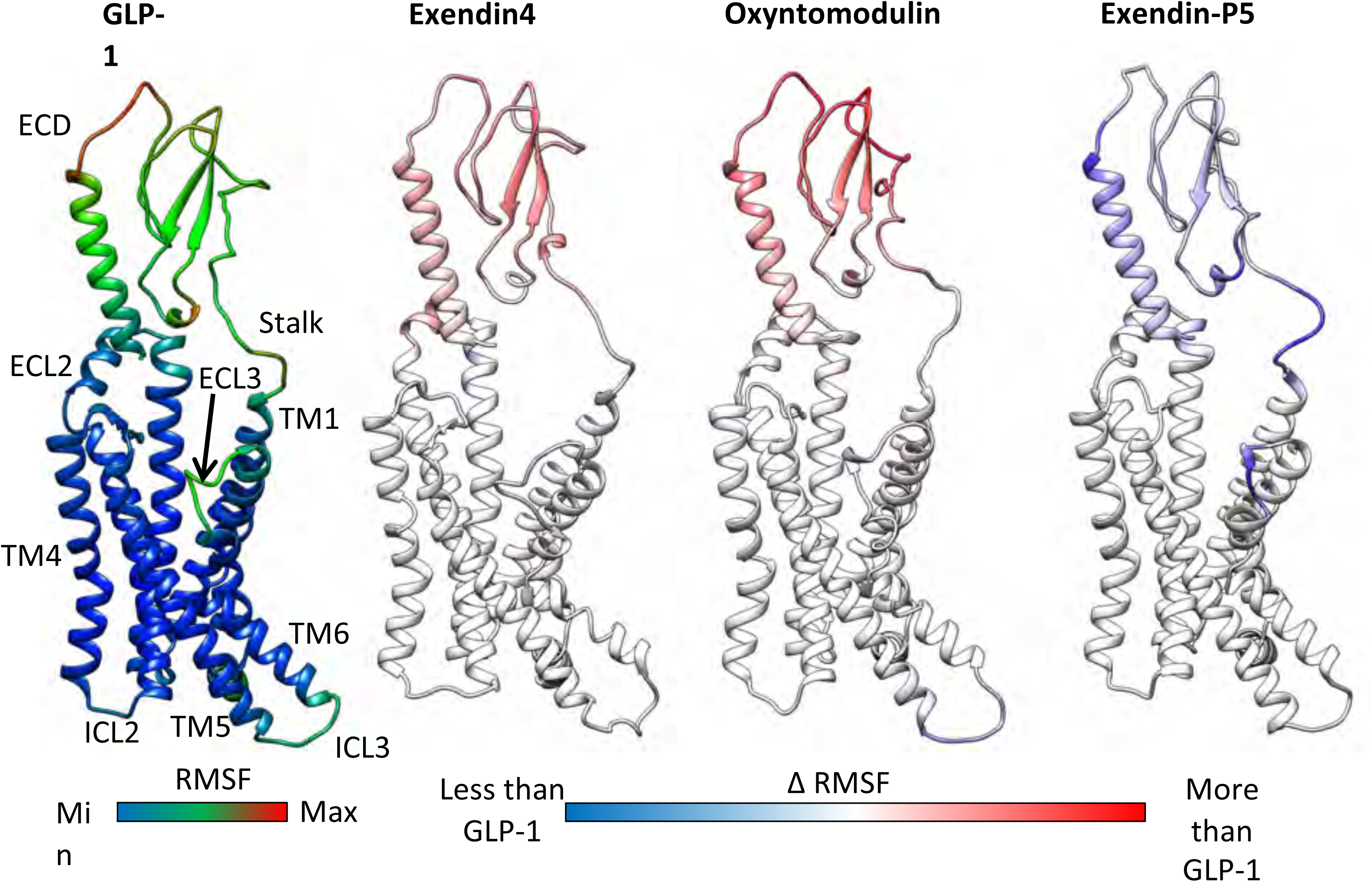
Comparison of the GLP1-R RMSF observed in MD simulations between the different GLP-1R:agonist complexes. For each residue, the RMSF in the presence of GLP-1 (left) is subtracted from the RMSF computed in the presence of the other agonists. Values are plotted on the backbone (ribbon) of the receptor.

**Supplemental Figure 13.**
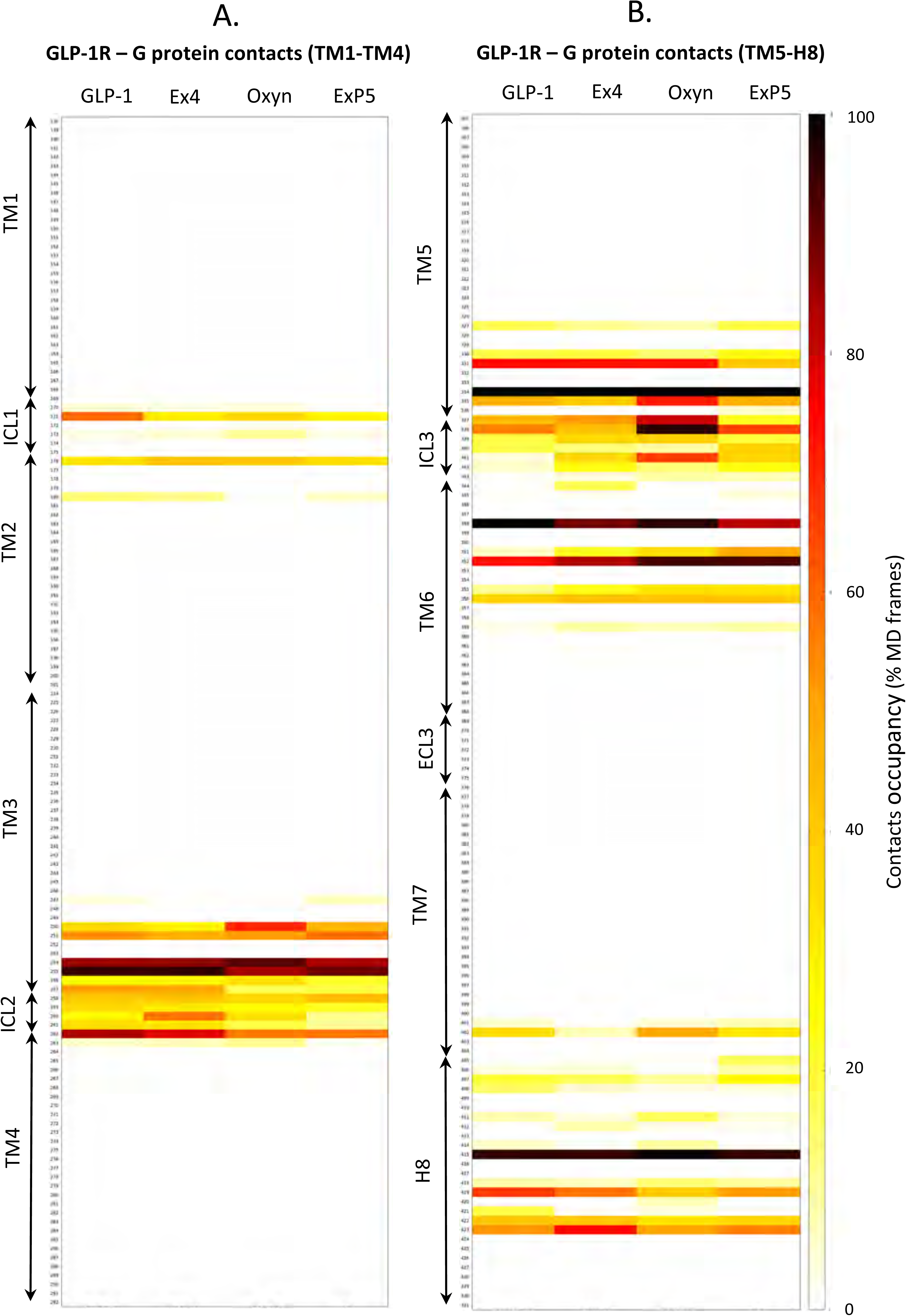
Heatmap showing the GLP-1R contacts with G_s_ in the presence of GLP-1, exendin-4, oxyntomodulin and exendin-P5. The GLP-1R primary sequence is divided into two segments: TM1 to TM4 (A), and TM5 to H8 (B). ECD, ECL1, and ECL2 have been omitted as these do not contact G_s_. For each agonist (GLP-1, exendin-4 (Ex4), oxyntomodulin (Oxyn), exendin-P5 (ExP5)), data are reported as the percentage of MD frames (occupancy) with at least one interatomic contact and normalized for the maximum occupancy.

**Supplemental Table 1.** Interactions between GLP-1R and GLP-1 (Zhang, et al. 2020), oxyntomodulin, exendin-4 and ExP5 (Liang, et al. 2018). Receptor TMD residues within 4 Å of the exendin-4 and oxyntomodulin within the static cryo-EM structures are detailed.

| GLP-1R:GLP-1:Gs |  | GLP-1R:Oxy:Gs |  | GLP-1R:Ex4:Gs |  | GLP-1R:ExP5:Gs |  |
| --- | --- | --- | --- | --- | --- | --- | --- |
| GLP-1R | GLP-1 | GLP-1R | Oxy | GLP-1R | Ex4 | GLP-1R | ExP5 |
| V237<sup>3.40</sup> | H7 | V237<sup>3.40</sup> | H1 | V237<sup>3.40</sup> | H1 | V237<sup>3.40</sup> | L2 |
| W306<sup>5.36</sup> | H7 | W306<sup>5.36</sup> | H1 |  |  |  |  |
| I309<sup>5.39</sup> | H7 |  |  |  |  |  |  |
| R310<sup>5.40</sup> | H7 (water mediated) | R310<sup>5.40</sup> | H1 |  |  | R310<sup>5.40</sup> | E1 |
| I313<sup>5.43</sup> | H7 |  |  |  |  | I313<sup>5.43</sup> | L2 |
| Q234<sup>3.37</sup> | H7 | Q234<sup>3.37</sup> | H1 |  |  |  |  |
|  |  | Y241<sup>3.44</sup> | H1 |  |  | A368<sup>6.57</sup> | E1 |
| Y241<sup>3.44</sup> | A8 (water mediated) |  |  |  |  |  |  |
| L384<sup>7.39</sup> | A8 | L384<sup>7.39</sup> | S2 |  |  | L384<sup>7.39</sup> | V3 |
| E387<sup>7.42</sup> | A8 | E387<sup>7.42</sup> | S2 |  |  | E387<sup>7.42</sup> | V3 |
| L388<sup>7.43</sup> | A8 | L388<sup>7.43</sup> | S2 |  |  | L388<sup>7.43</sup> | V3 |
| Y152<sup>1.47</sup> | E9 | Y152<sup>1.47</sup> | Q3 | Y152<sup>1.47</sup> | E3 | Y152<sup>1.47</sup> | D4 |
| R190<sup>2.60</sup> | E9 | V194<sup>2.64</sup> | Q3 | R190<sup>2.60</sup> | E3 | V194<sup>2.64</sup> | D4 |
| Y241<sup>3.44</sup> | E9 (water mediated) | Y148<sup>1.43</sup> | Q3 | Y148<sup>1.43</sup> | E3 | K197<sup>2.67</sup> | D4 |
| L388<sup>7.43</sup> | E9 (water mediated) | L388<sup>7.43</sup> | Q3 |  |  | M233<sup>3.36</sup> | D4 |
| T298<sup>ECL2</sup> | G10 (water mediated) |  |  |  |  |  |  |
| N300<sup>ECL2</sup> | G10 |  |  |  |  |  |  |
| W306<sup>5.36</sup> | G10 | W306<sup>5.36</sup> | G4 | W306<sup>5.36</sup> | G4 |  |  |
| W306<sup>5.36</sup> | T11 | W306<sup>5.36</sup> | T5 |  |  | W306<sup>5.36</sup> | N5 |
| D372<sup>ECL3</sup> | T11 |  |  |  |  | Q234<sup>3.37</sup> | N5 |
| R380<sup>7.35</sup> | T11 |  |  |  |  |  |  |
| L384<sup>7.39</sup> | T11 |  |  |  |  |  |  |
| L141<sup>1.36</sup> | F12 | L141<sup>1.36</sup> | F6 | L141<sup>1.36</sup> | F6 |  |  |
| L144<sup>1.39</sup> | F12 | L144<sup>1.39</sup> | F6 | L144<sup>1.39</sup> | F6 |  |  |
| Y148<sup>1.43</sup> | F12 |  |  |  |  | L384<sup>7.39</sup> | V7 |
| L388<sup>7.43</sup> | F12 |  |  |  |  | L388<sup>7.43</sup> | V7 |
| K197<sup>2.67</sup> | T13 | K197<sup>2.67</sup> | T7 | K197<sup>2.67</sup> | T7 |  |  |
| F230<sup>3.33</sup> | T13 |  |  |  |  |  |  |
| M233<sup>3.36</sup> | T13 |  |  | M233<sup>3.36</sup> | T7 |  |  |
| T298<sup>ECL2</sup> | T13 | T298<sup>ECL2</sup> | T7 |  |  |  |  |
| T298<sup>ECL2</sup> | S14 | T298<sup>ECL2</sup> | S8 | T298<sup>ECL2</sup> | S8 |  |  |
| R299<sup>ECL2</sup> | S14 |  |  |  |  |  |  |
| N300<sup>ECL2</sup> | S14 | N300<sup>ECL2</sup> | S8 |  |  |  |  |
| R380<sup>7.35</sup> | D15 | R380<sup>7.35</sup> | D9 |  |  | R380<sup>7.35</sup> | D10 |
| L384<sup>7.39</sup> | D15 |  |  |  |  |  |  |
| L201<sup>2.71</sup> | V16 | L201<sup>2.71</sup> | Y10 |  |  | L141<sup>1.36</sup> | L11 |
| Y205<sup>ECL1</sup> | S17 |  |  |  |  | L201<sup>2.71</sup> | S12 |
| T298<sup>ECL2</sup> | S17 | T298<sup>ECL2</sup> | S11 | T298<sup>ECL2</sup> | S11 | R298<sup>ECL2</sup> | S12 |
| R299<sup>ECL2</sup> | S17 | R299<sup>ECL2</sup> | S11 | R299<sup>ECL2</sup> | S11 |  |  |
| R299<sup>ECL2</sup> | S18 | R299<sup>ECL2</sup> | K12 |  |  | R299<sup>ECL2</sup> | K13 |
| P137<sup>1.32</sup> | Y19 |  |  |  |  |  |  |
| E138<sup>1.33</sup> | Y19 | E138<sup>1.33</sup> | Y13 |  |  | E138<sup>1.33</sup> | Q14 |
|  |  | L141<sup>1.36</sup> | Y13 |  |  | L141<sup>1.36</sup> | Q14 |
| Y205<sup>ECL1</sup> | L20 | Y205<sup>ECL1</sup> | L14 |  |  | Y205<sup>ECL1</sup> | M15 |
|  |  |  |  |  |  | L201<sup>2.71</sup> | M15 |
|  |  |  |  |  |  | K202<sup>2.72</sup> | M15 |
|  |  |  |  |  |  | S206<sup>ECL1</sup> | M15 |
| V30<sup>ECD</sup> | E21 |  |  |  |  |  |  |
| S31<sup>ECD</sup> | E21 |  |  |  |  |  |  |
| L32<sup>ECD</sup> | E21 |  |  |  |  | L32<sup>ECD</sup> | E16 |
| Y205<sup>ECL1</sup> | E21 | Y205<sup>ECL1</sup> | D15 | Y205<sup>ECL1</sup> | E15 | Y205<sup>ECL1</sup> | E16 |
| Q221<sup>ECL1</sup> | E21 |  |  |  |  |  |  |
| R299<sup>ECL2</sup> | E21 | R299<sup>ECL2</sup> | D15 |  |  | R299<sup>ECL2</sup> | E16 |
| L32<sup>ECD</sup> | A24 |  |  |  |  |  |  |
| Q210<sup>ECL1</sup> | A24 | Y205<sup>ECL1</sup> | R18 |  |  | Y205<sup>ECL1</sup> | A19 |
|  |  |  |  |  |  | L32<sup>ECD</sup> | V20 |
|  |  |  |  |  |  | P90<sup>ECD</sup> | V20 |
| E128<sup>ECD</sup> | K26 |  |  |  |  |  |  |
| Q210<sup>ECL1</sup> | E27 |  |  |  |  |  |  |
| L32<sup>ECD</sup> | F28 |  |  |  |  |  |  |
| T35<sup>ECD</sup> | F28 |  |  |  |  |  |  |
| W39<sup>ECD</sup> | F28 |  |  |  |  | W39<sup>ECD</sup> | F23 |
| W214<sup>ECL1</sup> | F28 |  |  | W214<sup>ECL1</sup> | F22 |  | F23 |
| T35<sup>ECD</sup> | I29 |  |  |  |  | V20<sup>ECD</sup> | I24 |
| Y69<sup>ECD</sup> | I29 |  |  |  |  | Y69<sup>ECD</sup> | I24 |
| P90<sup>ECD</sup> | I29 |  |  |  |  | P90<sup>ECD</sup> | I24 |
| W91<sup>ECD</sup> | I29 |  |  |  |  | W91<sup>ECD</sup> | I24 |
| W214<sup>ECL1</sup> | W31 |  |  | W214<sup>ECL1</sup> | W25 | W214<sup>ECL1</sup> | W26 |
|  |  |  |  |  |  | H212<sup>ECL1</sup> | W26 |
| W39<sup>ECD</sup> | L32 |  |  |  |  |  |  |
| E68<sup>ECD</sup> | L32 |  |  |  |  |  |  |
| Y69<sup>ECD</sup> | L32 |  |  |  |  |  |  |
| Y69<sup>ECD</sup> | V33 |  |  |  |  |  |  |
| R121<sup>ECD</sup> | V33 |  |  |  |  |  |  |
| W39<sup>ECD</sup> | R36 |  |  |  |  |  |  |
| E68<sup>ECD</sup> | R36 |  |  |  |  |  |  |

**Supplementary Table 2.**
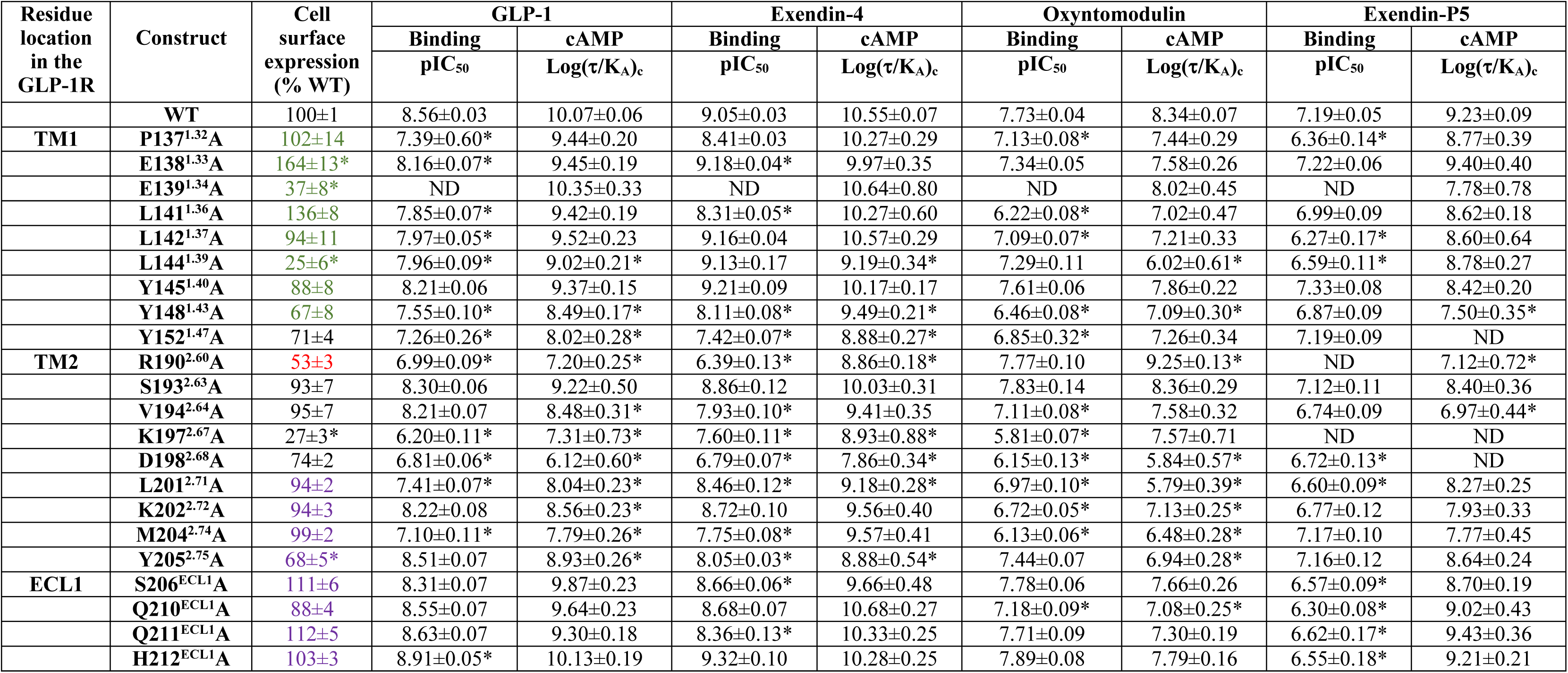
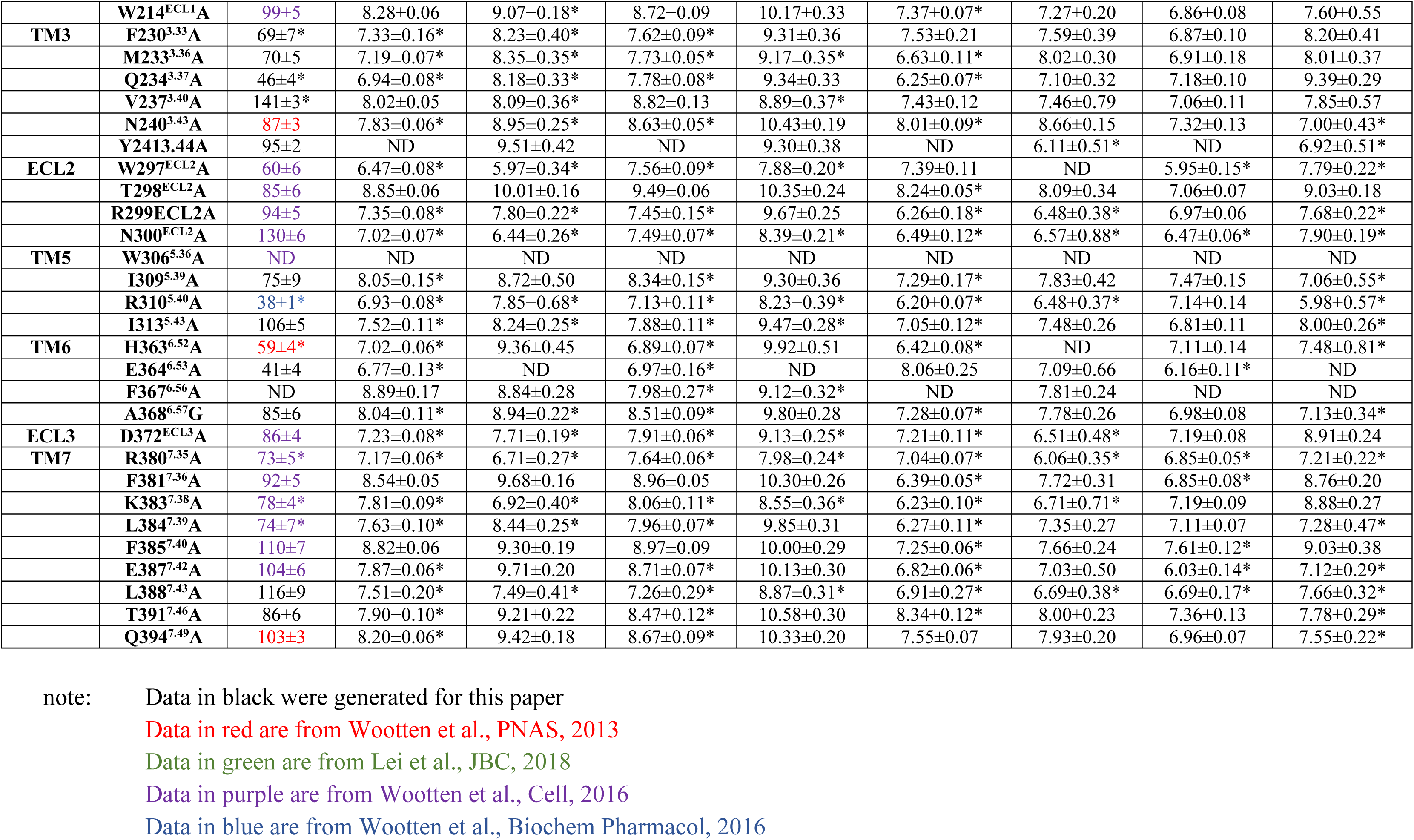
The effects of TMD mutations on peptide affinity and cAMP signalling efficiency. Mutant and WT GLP-1Rs were stably expressed in ChoFlpIn cells and competition inhibition binding curves **(Supplementary Figures 5-6)** and cAMP accumulation concentration response curves (Supplementary Figures 7-8) were generated for each construct for the four agonists. Binding data were analyzed using a three-parameter logistic equation to determine pIC_50_ values, which represent the negative logarithm of the concentration of ligand that inhibits binding of half the total concentration of radiolabelled antagonist, ^125^I-exendin(9-39). Cell surface expression was determined through antibody detection of the N-terminal c-myc epitope label, with data expressed as a maximum of wildtype human GLP-1R expression. cAMP data were analysed with an operational model of agonism to determine logτ/K_A_ values, which were then corrected to cell surface expression data. Values are expressed as mean ± s.e.m. Data were analysed with one-way analysis of variance and Dunnett’s post test (* p < 0.05). ND means data that were unable to be experimentally defined.

**Supplementary Table 3.**
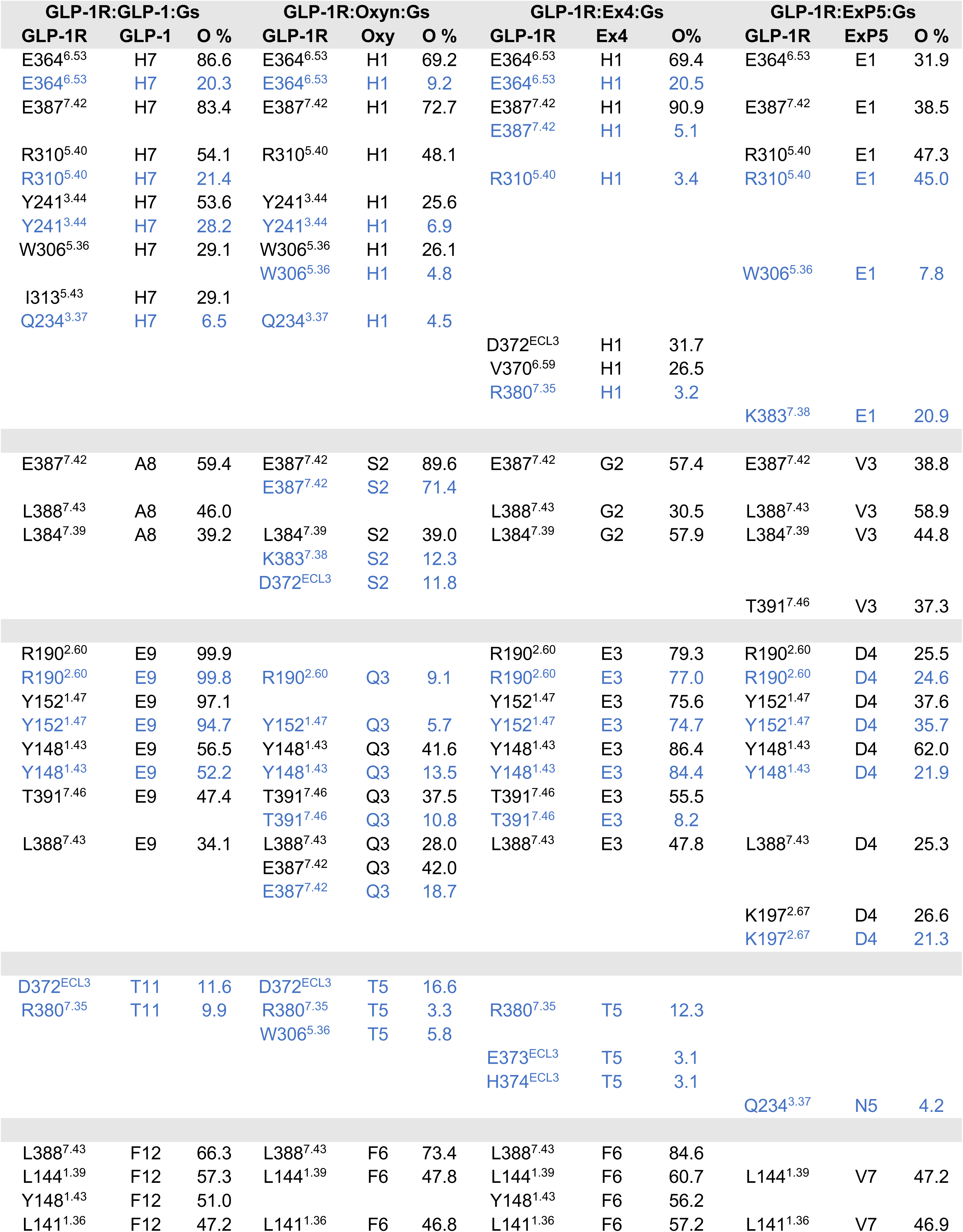
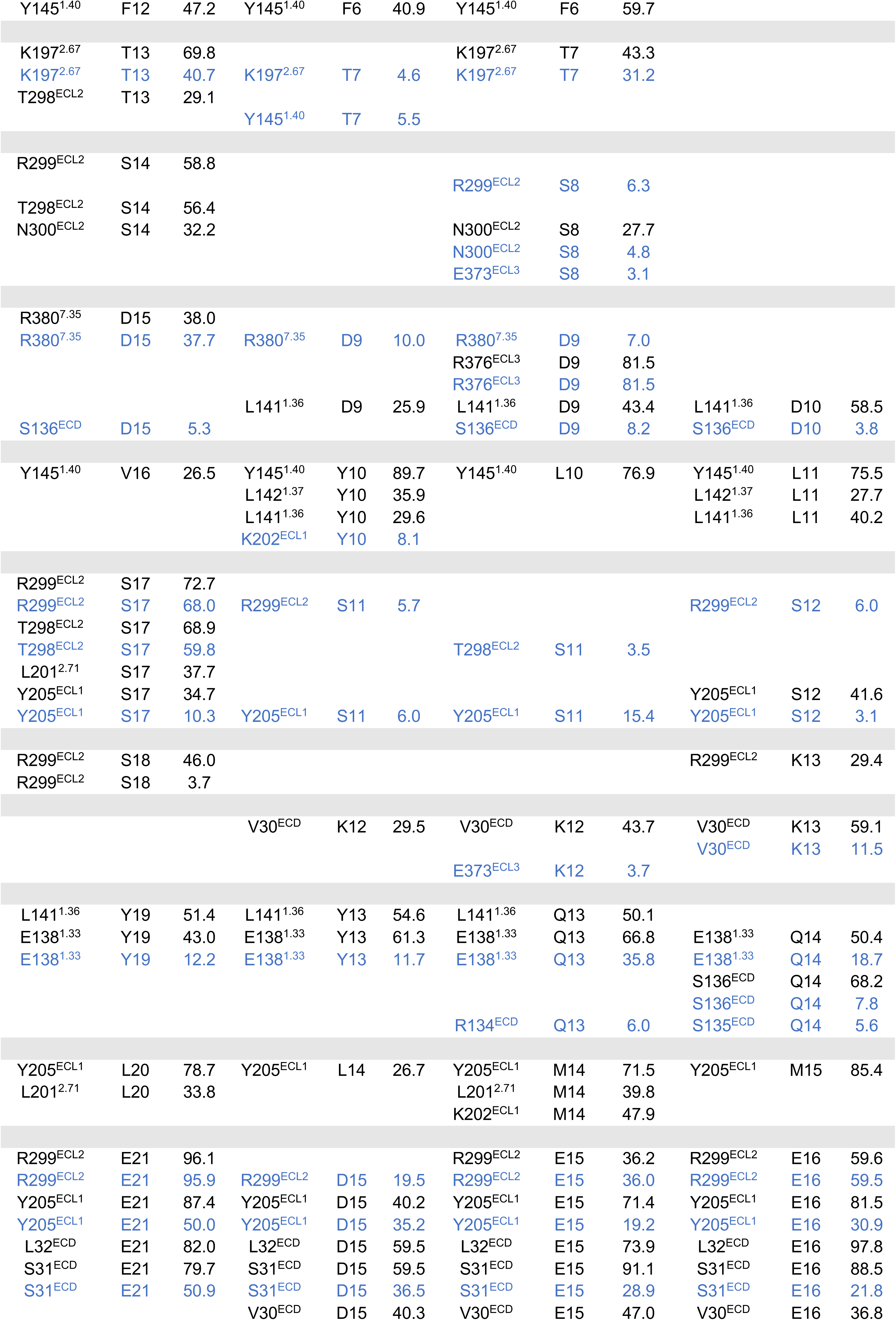
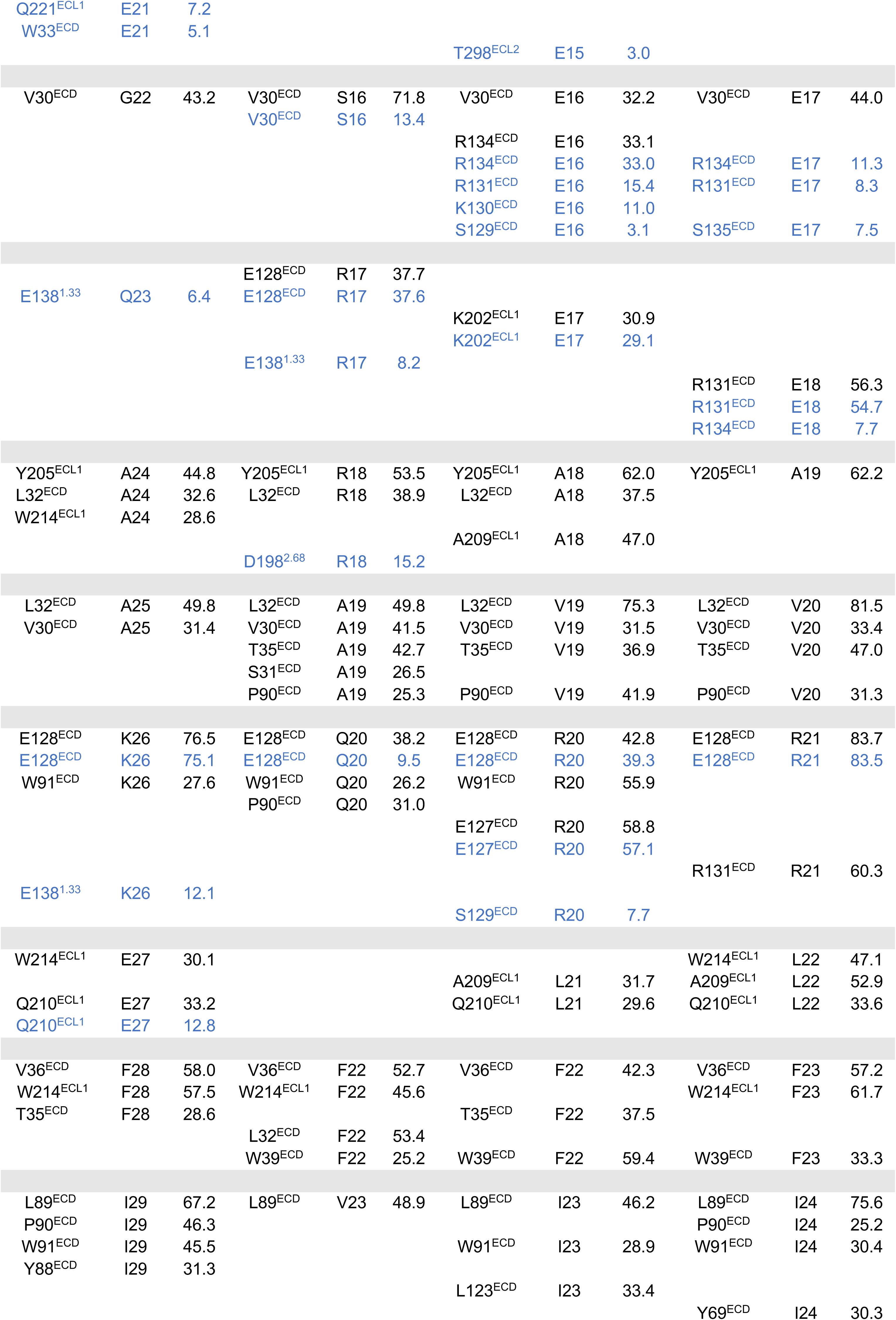
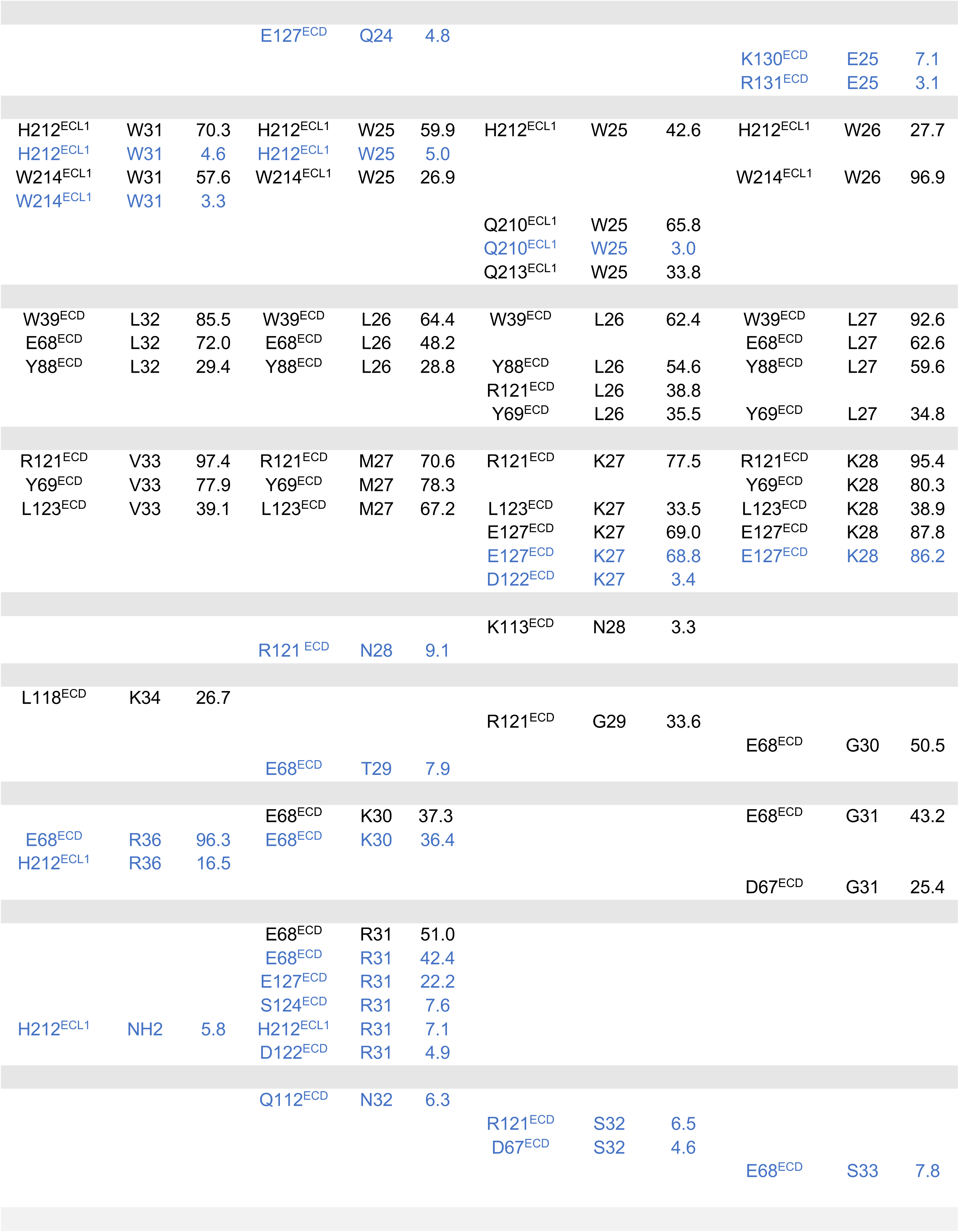
Main contacts between GLP-1R and GLP-1, exendin-4 (Ex4), oxyntomodulin (Oxyn), and exendin-P5 (ExP5), during MD simulations. Data are expressed as the occupancy (% of frames) in which the interactions were present. Black = all contacts (<25% occupancy not shown), blue = Main hydrogen bonds (side chain-side chain or side chain-backbone). Interactions are reported in order of the peptide sequence for each peptide (NB – some weaker and more transient contacts observed in the MD are not shown).

**Supplementary Table 4.**
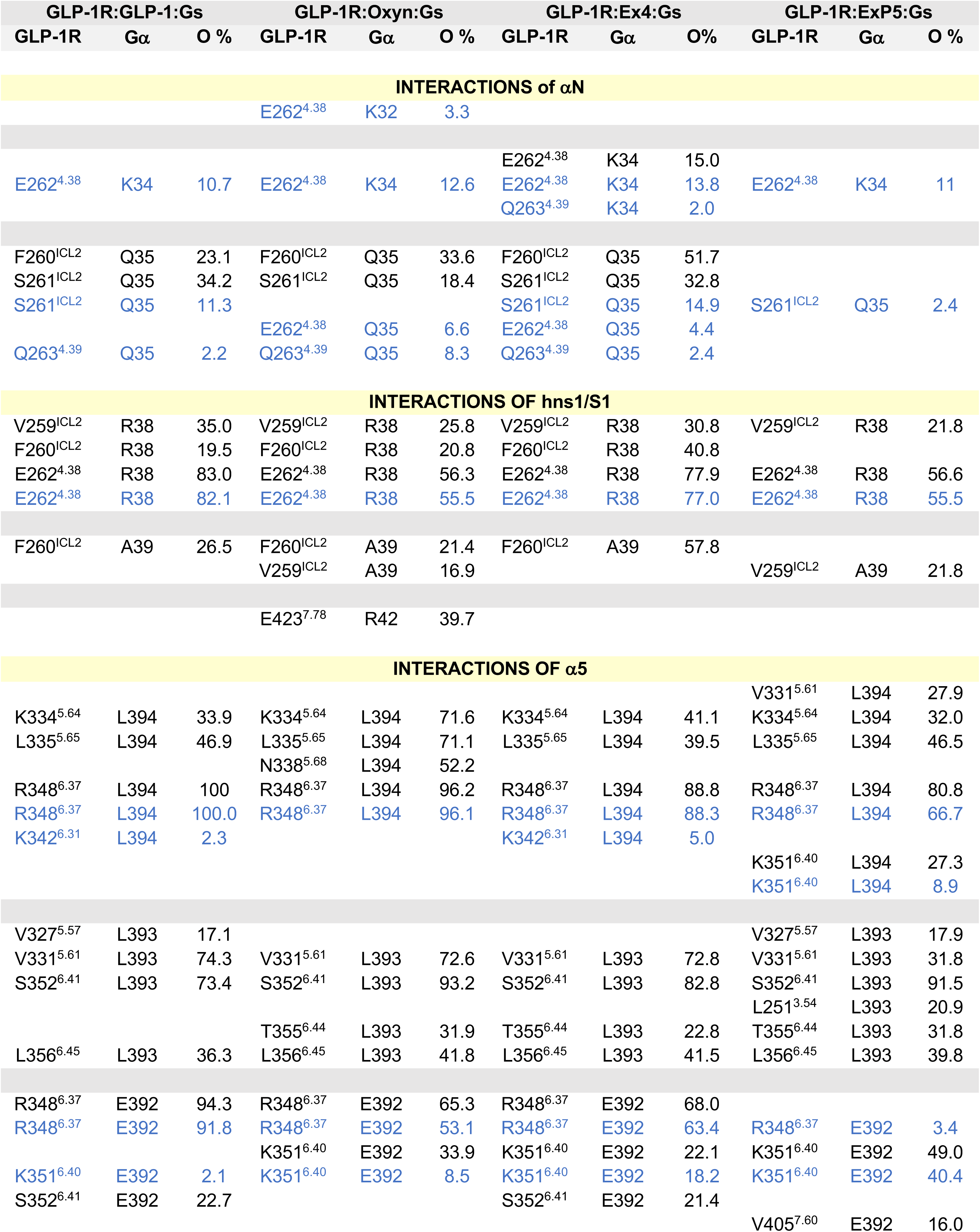
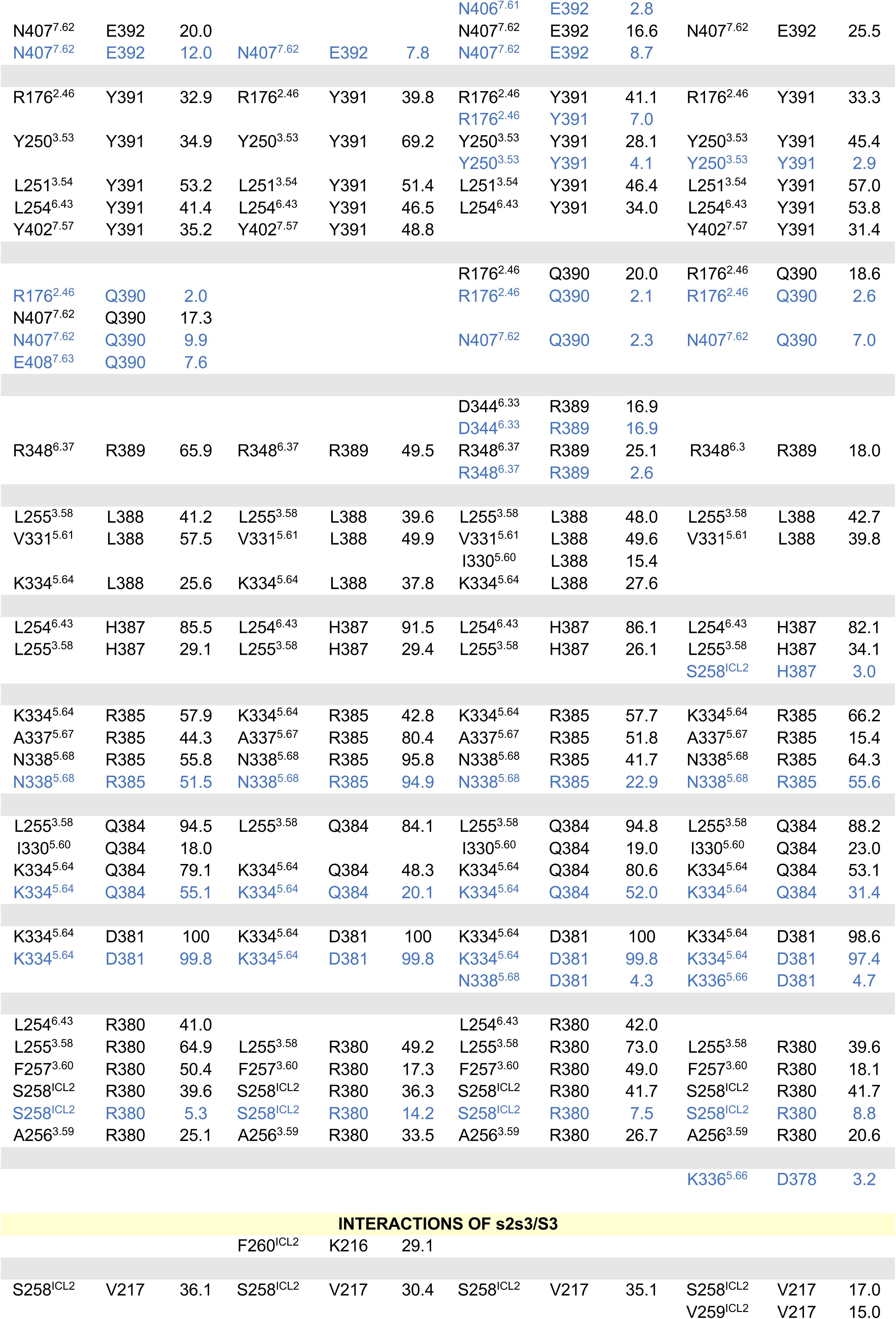
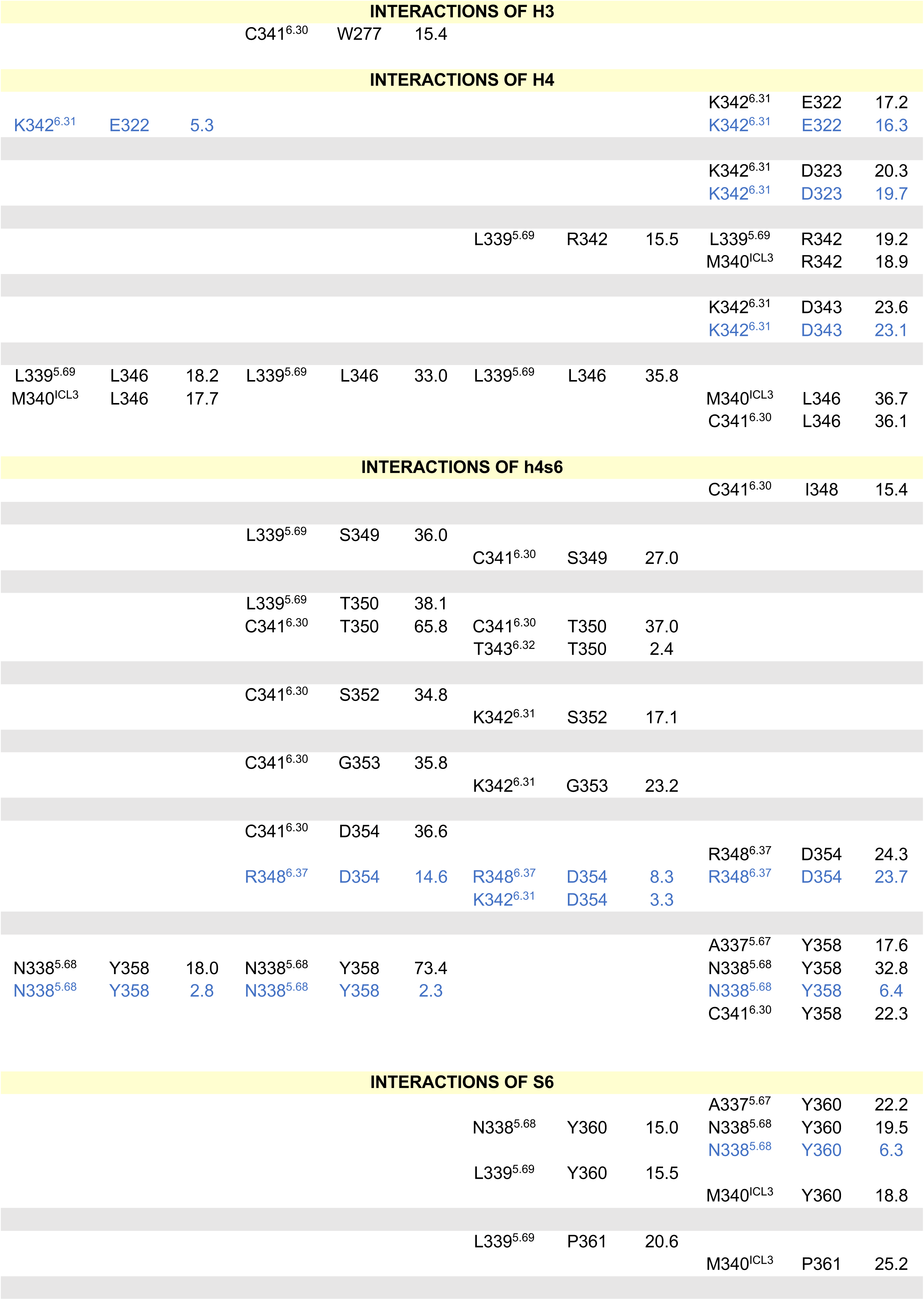
Main contacts between GLP-1R (in complex with GLP-1, Ex4, Oxyn, and ExP5) and the G protein G*α* subunit, during MD simulations. Data are expressed as the occupancy (% of frames) in which the interactions were present. Black = all contacts (<15% occupancy not shown), blue = main hydrogen bonds (side chain-side chain or side chain-backbone - (NB – some weaker and more transient contacts observed in the MD are not shown). Interactions are listed by regions within the G protein (using the CGN nomenclature).

**Supplementary Table 5.** Main contacts between GLP-1R (in complex with GLP-1, Ex4, Oxyn, and ExP5) and the G protein G*β* subunit, during MD simulations. Data are expressed as the occupancy (% of frames) in which the interactions were present. Black = all contacts (<15% occupancy not shown), blue = main hydrogen bonds (side chain-side chain or side chain-backbone) - (NB – some weaker and more transient contacts observed in the MD are not shown). Interactions are listed in order of the sequence of the G*β* subunit.

| GLP-1R:GLP-1:Gs |  |  | GLP-1R:Oxy:Gs |  |  | GLP-1R: Ex4:Gs |  |  | GLP-1R:ExP5:Gs |  |  |
| --- | --- | --- | --- | --- | --- | --- | --- | --- | --- | --- | --- |
| GLP-1R | Gβ | O % | GLP-1R | Gβ | O % | GLP-1R | Gβ | O % | GLP-1R | Gβ | O % |
| E423<sup>7.78</sup> | R42 | 52.5 | E423<sup>7.78</sup> | R42 | 39.7 | E423<sup>7.78</sup> | R42 | 25.4 | E423<sup>7.78</sup> | R42 | 55.1 |
| E423<sup>7.78</sup> | R42 | 40.1 | E423<sup>7.78</sup> | R42 | 24.7 | E423<sup>7.78</sup> | R42 | 16.8 | E423<sup>7.78</sup> | R42 | 44.8 |
| E423<sup>7.78</sup> | Q44 | 18.4 |  |  |  |  |  |  |  |  |  |
| E423<sup>7.78</sup> | Q44 | 7.5 | E423<sup>7.78</sup> | Q44 | 2.7 | E423<sup>7.78</sup> | Q44 | 2.7 |  |  |  |
|  |  |  | L422<sup>7.77</sup> | R46 | 15.2 |  |  |  |  |  |  |
| E423<sup>7.78</sup> | R46 | 48.4 | E423<sup>7.78</sup> | R46 | 49.7 | E423<sup>7.78</sup> | R46 | 74.2 | E423<sup>7.78</sup> | R46 | 33.1 |
| E423<sup>7.78</sup> | R46 | 47.3 | E423<sup>7.78</sup> | R46 | 41.0 | E423<sup>7.78</sup> | R46 | 61.4 | E423<sup>7.78</sup> | R46 | 29.0 |
|  |  |  | H171<sup>ICL1</sup> | R52 | 35.9 | H171<sup>ICL1</sup> | R52 | 30.0 |  |  |  |
|  |  |  | H171<sup>ICL1</sup> | R52 | 4.7 | H171<sup>ICL1</sup> | R52 | 10.2 |  |  |  |
|  |  |  | E412<sup>7.67</sup> | R52 | 2.1 | E412<sup>7.67</sup> | R52 | 8.5 |  |  |  |
| R421<sup>7.76</sup> | D291 | 17.7 |  |  |  |  |  |  |  |  |  |
| R421<sup>7.76</sup> | D291 | 17.7 |  |  |  |  |  |  |  |  |  |
| R414<sup>7.69</sup> | D291 | 5.4 |  |  |  |  |  |  |  |  |  |
|  |  |  | E418<sup>7.73</sup> | N293 | 9.6 |  |  |  | E418<sup>7.73</sup> | N293 | 7.8 |
| L422<sup>7.77</sup> | N293 | 30.2 |  |  |  | L422<sup>7.77</sup> | N293 | 19.9 | L422<sup>7.77</sup> | N293 | 16.6 |
| E423<sup>7.78</sup> | R304 | 19.0 | E423<sup>7.78</sup> | R304 | 18.9 |  |  |  |  |  |  |
| E423<sup>7.78</sup> | R304 | 14.3 | E423<sup>7.78</sup> | R304 | 18.5 | E423<sup>7.78</sup> | R304 | 4.6 | E423<sup>7.78</sup> | R304 | 10.8 |
|  |  |  |  |  |  |  |  |  | E423<sup>7.78</sup> | G306 | 22.4 |
| L422<sup>7.77</sup> | V307 | 40.8 | L422<sup>7.77</sup> | V307 | 32.0 | L422<sup>7.77</sup> | V307 | 39.2 | L422<sup>7.77</sup> | V307 | 35.9 |
| E423<sup>7.78</sup> | V307 | 26.4 | E423<sup>7.78</sup> | V307 | 20.8 |  |  |  | E423<sup>7.78</sup> | V307 | 31.6 |
|  |  |  | R419<sup>7.74</sup> | A309 | 16.4 | R419<sup>7.74</sup> | A309 | 30.3 | R419<sup>7.74</sup> | A309 | 16.2 |
| L422<sup>7.77</sup> | A309 | 37.7 | L422<sup>7.77</sup> | A309 | 23.7 | L422<sup>7.77</sup> | A309 | 41.4 | L422<sup>7.77</sup> | A309 | 32.0 |
|  |  |  | K415<sup>7.70</sup> | G310 | 29.5 |  |  |  |  |  |  |
| R419<sup>7.74</sup> | G310 | 49.4 | R419<sup>7.74</sup> | G310 | 38.0 | R419<sup>7.74</sup> | G310 | 49.6 | R419<sup>7.74</sup> | G310 | 42.8 |
| R419<sup>7.74</sup> | H311 | 63.5 | R419<sup>7.74</sup> | H311 | 33.3 | R419<sup>7.74</sup> | H311 | 39.9 | R419<sup>7.74</sup> | H311 | 45.4 |
| H171<sup>ICL1</sup> | D312 | 59.9 | H171<sup>ICL1</sup> | D312 | 16.5 |  |  |  | H171<sup>ICL1</sup> | D312 | 27.2 |
| H171<sup>ICL1</sup> | D312 | 34.6 | H171<sup>ICL1</sup> | D312 | 4.2 |  |  |  | H171<sup>ICL1</sup> | D312 | 22.5 |
| K415<sup>7.70</sup> | D312 | 95.1 | K415<sup>7.70</sup> | D312 | 99.1 | K415<sup>7.70</sup> | D312 | 94.8 | K415<sup>7.70</sup> | D312 | 93.9 |
| K415<sup>7.70</sup> | D312 | 92.6 | K415<sup>7.70</sup> | D312 | 98.9 | K415<sup>7.70</sup> | D312 | 89.7 | K415<sup>7.70</sup> | D312 | 91.3 |
| R419<sup>7.74</sup> | D312 | 66.8 | R419<sup>7.74</sup> | D312 | 39.4 | R419<sup>7.74</sup> | D312 | 57.9 | R419<sup>7.74</sup> | D312 | 50.5 |
| R419<sup>7.74</sup> | D312 | 66.4 | R419<sup>7.74</sup> | D312 | 39.3 | R419<sup>7.74</sup> | D312 | 57.5 | R419<sup>7.74</sup> | D312 | 50.2 |

**Supplemental Table 6:** CryoEM Data collection and modelling refinement statistics.

| **Data Collection** | GLP-1R:DNGs:EX4 | GLP-1R:DNGs:Oxyntomodullin |
| --- | --- | --- |
| Micrographs | 8816 | 3181 |
| Electron dose (e<sup>-</sup>/Å<sup>2</sup>) | 48 | 50 |
| Voltage (kV) | 300 | 300 |
| Pixel size (Å) | 0.87 | 1.06 |
| Defocus range (μm) | 0.5-1.5 | 0.5 |
| Symmetry imposed | C1 | C1 |
| Particles (final map) | 277,494 | 366,351 |
| Resolution (0.143 FSC)(Å) | 3.7 | 3.3 |
| **Refinement** |  |  |
| CC<sub>map_model</sub> |  |  |
| Map sharpening B factor (Å<sup>2</sup>) | -137 | -50 |
| **Model Quality** |  |  |
| R.m.s. deviations |  |  |
| Bond length (Å) | 0.006 | 0.002 |
| Bond angles (°) | 0.890 | 0.447 |
| Ramachandran |  |  |
| Favoured (%) | 92.98 | 97.95 |
| Outliers (%) | 0 | 0 |
| Rotamer outliers | 0.1 | 0.68 |
| C-Beta deviations (%) | 0 | 0 |
| Clashscore | 8.28 | 3.05 |
| MolProbity score | 1.90 | 1.11 |

**Supplemental Video 1. MD simulations of the peptide bound GLP-1R:G_s_ complexes.** Comparison of the dynamic (merged MD replicas) of the four GLP-1R:agonist:G_s_ complexes. GLP-1R is shown as a purple ribbon. The residues forming intermolecular contacts and hydrogen bonds (transient red dotted lines) are shown as sticks. GLP-1, exendin-4 (Ex4), oxyntomodulin (Oxyn), and exendin-P5 (ExP5) are shown in red, green, yellow, and cyan respectively.

## Notes

### Competing Interest Statement

The authors have declared no competing interest.

